# Expression Atlas Of *Dmrt* Genes Across Sex And Development – Functional Insights From The Mouse Olfactory System

**DOI:** 10.1101/2025.09.11.674830

**Authors:** Rafael Casado-Navarro, Ana Bermejo-Santos, Rodrigo Torrillas-de la Cal, María Pilar Madrigal, Virgilia Olivé, Li Ying Chen-Chen, Sonia Amorós-Bru, Sandra Jurado, Esther Serrano-Saiz

## Abstract

The ancient DMRT family of transcription factors has been proposed as evolutionarily conserved effectors of sexual differentiation. While brain sexual differentiation has traditionally been attributed to the sex chromosome complement (XX or XY) and steroid sex hormones, the downstream effector mechanisms controlled by these factors remain elusive. To elucidate the role of *Dmrts* in the mammalian brain sexual differentiation, we generated a comprehensive expression atlas for all family members (*Dmrt1-7*) in the mouse brain. We used *in situ* hybridization to examine both sexes across various developmental stages. Our findings revealed that all *Dmrts*, except *Dmrt7*, are expressed in the brain. This study expands our understanding of the DMA-*Dmrt* subfamily beyond pallial structures and identifies their expression maintenance in adult neurogenic sites. For the first time, we described the neuronal expression of *Dmrt2* and *Dmrt6*. Mouse *Dmrts* did not exhibit clear sexually dimorphic patterns but showed quantitative differences in expression levels between the sexes. We demonstrated that most *Dmrts* are maintained in postmitotic neurons during both embryonic and postnatal stages, suggesting potential interactions with steroid hormones during organizational and activational phases.

As proof of concept, our comprehensive analysis of *Dmrt5* expression revealed its prominent presence in the mouse olfactory system, which is fundamental for controlling sex-specific innate behaviors. The absence of *Dmrt5* affects the main olfactory epithelium, where sensory neurons are located; however, mis-patterning phenotypes observed in the olfactory bulb and the piriform cortex distinctly affect male and female embryos, revealing the interaction of *Dmrt5* with sex in deeper integrative layers of innate neural circuits.

Our results provide a valuable resource for uncovering novel sites and mechanisms of sexual differentiation in the mammalian nervous system, potentially contributing to the sex bias observed in the prevalence and symptomatology of psychiatric disorders.

## INTRODUCTION

Across phylogeny, the nervous system (NS) presents sex differences that are responsible for sex-specific behaviors. In humans, brain sexual variations might uncover sex-related susceptibilities to mental disorders^1^. Sex hormones and genetic factors regulate the molecular mechanisms underlying the specification and maintenance of NS sex differences^2^. Among them, the ancient family of *Dmrt* genes (from *<u>d</u>oublesex* and *<u>m</u>ab-3* related transcription factors) emerged as conserved effectors in the development of sex-specific traits of different animal species^3–5^. Among all the *Dmrt*s*, Dmrt1* is the only one with a conserved role in the gonadal sex determination across vertebrates. Moreover, in mice, *Dmrt1* is essential for male somatic and germ cell differentiation, and in humans, deletions and point mutations in *DMRT1* are associated with 46XY complete gonadal dysgenesis^6^.

The role of *Dmrts* in the sexual differentiation of the NS has been primarily studied in invertebrates such as *Drosophila melanogaster* and *Caenorhabditis elegans*. In these organisms, species-specific sex determination cues control a sexually dimorphic expression pattern for *Dmrts* in a binary mode (reviewed in^5^). The sex-specific expression of *Dmrts* in one sex versus the other generates dimorphic neuronal circuits from which sex-specific behaviors emerge. To accomplish this, *Dmrt*s utilize distinct mechanisms, involving sex-specific control of neuron number, neuronal identity, and synaptic connectivity. Sex differences in neuron number arise through sex-specific *dsx* isoforms that promote proliferation in males or cell death in females in *Drosophila*^7–10^. Regulation of sex-specific neuronal identity is a common feature among *Dmrts* in *C. elegans*. For instance, *mab-3* promotes the neurogenesis of male-specific sensory ray neurons^11–13^, while in these neurons, *dmd-3* and *mab-23* determine their neurotransmitter identities^14,15^. Regarding sexually dimorphic connectivity, the female-specific isoform of *dsx* represses midline crossing of axons from foreleg gustatory receptor neurons^16^, and *dmd-3* orchestrates the drastic male wiring remodeling of the sex-shared PHC neurons^17^. Most animal genomes contain multiple *Dmrts* defined by a common DNA-binding domain known as the DM domain. In mammals, there are seven *Dmrts*, *Dmrt1* to *Dmrt7*^18–20^, and up to three *Dmrt-like* genes (*Dmrt8.1, Dmrt8.2,* and *Dmrt8.3*) clustered in the X chromosome, which lack a functional DM domain^21^. However, the DM domain-encoding sequence of *Dmrt8* is not conserved in the mouse genome and, therefore, will not be considered in this study. *Dmrt* family genes exhibit limited sequence homology outside the highly conserved DM domain, except for a subgroup that contains an additional domain, the DMA domain. This subfamily includes *Dmrt3*, *Dmrt4* (also known as *Dmrta1*), and *Dmrt5* (also known as *Dmrta2*) in mammals, *dmrt93B* and *dmrt99B* in *D. melanogaster*, and *dmd-4* in *C. elegans*. From this point forward, we will collectively refer to the *Dmrt3*-*5* genes as DMA-*Dmrts*.

A comprehensive characterization of *Dmrt* gene expression patterns in the mammalian NS is crucial for elucidating novel mechanisms of sexual differentiation. Previous work mainly focused on DMA-*Dmrt* expression at early developmental stages, mostly in the telencephalon, for which excellent reviews exist^19,22^. However, no systematic comparisons between the sexes were undertaken in such studies. Furthermore, no studies have addressed *Dmrt* expression at late embryonic and postnatal stages, when novel functions or interactions with sex hormones might occur. Also, available gene expression atlases, such as the Allen Developing Mouse Brain Atlas or EURExpress, show poor or absent *in situ* hybridization (ISH) signals for *Dmrts*.

In this work, we generated a comprehensive map across sex and development for all *Dmrts* found in the mammalian genome. The stages considered for this study span the period at the peak of neurogenesis and gonadal hormonal secretion, encompassing embryonic (E) days 12.5, 14.5, and 18.5, postnatal (P) day 7, and, in some cases, adulthood (P56). Our study is primarily based on ISH; however, other techniques were also employed, including quantitative RT-PCR (qRT-PCR) and immunofluorescence (IF) and 3D brain reconstructions. We provide a molecular atlas of individual expression of *Dmrt1*, *Dmrt2, Dmrt3, Dmrt4, Dmrt5,* and *Dmrt6* in male and female embryos and postnatal animals. *Dmrt7* was not detected in the mouse NS. Our work is an invaluable resource for exploring the function of the DMRT transcription factors (TFs) in the mammalian brain, including the development of sexual dimorphisms in the NS. Moreover, it will help identify markers that could serve as a resource for functional investigations.

## RESULTS

### All *Dmrt* genes, except *Dmrt7*, are detected in the embryonic NS

We initiated our analysis at E12.5, before testosterone can be secreted by the fetal testis (approximately at E13.0). We followed the E13.5 sagittal atlas of the Allen Developing Mouse Brain Atlas (ADMBA, available at: https://developingmouse.brain-map.org/static/atlas), which is based on the prosomeric model established by Luis Puelles^23^. To a lesser extent, we also used the Prenatal Mouse Brain Atlas at E12.5^24^, which includes coronal sections, while maintaining the nomenclature of the ADMBA atlas, except for the choroid plexus (ChPl), the piriform cortex (Pir), and the gigantocellular reticular nucleus (GRN). This analysis relied solely on anatomical landmarks. When precision was uncertain, we deliberately avoided overly detailed annotations and instead assigned expression to broader topological regions. In **Figure 1**, only a selection of images of one sex is depicted to complement **Table 1**, which compiles the expression of all *Dmrts* at E12.5. For full access to the expression data and sex comparisons, see **Repository link:** https://www.informatics.jax.org/reference/J:364871.

**Figure 1.**
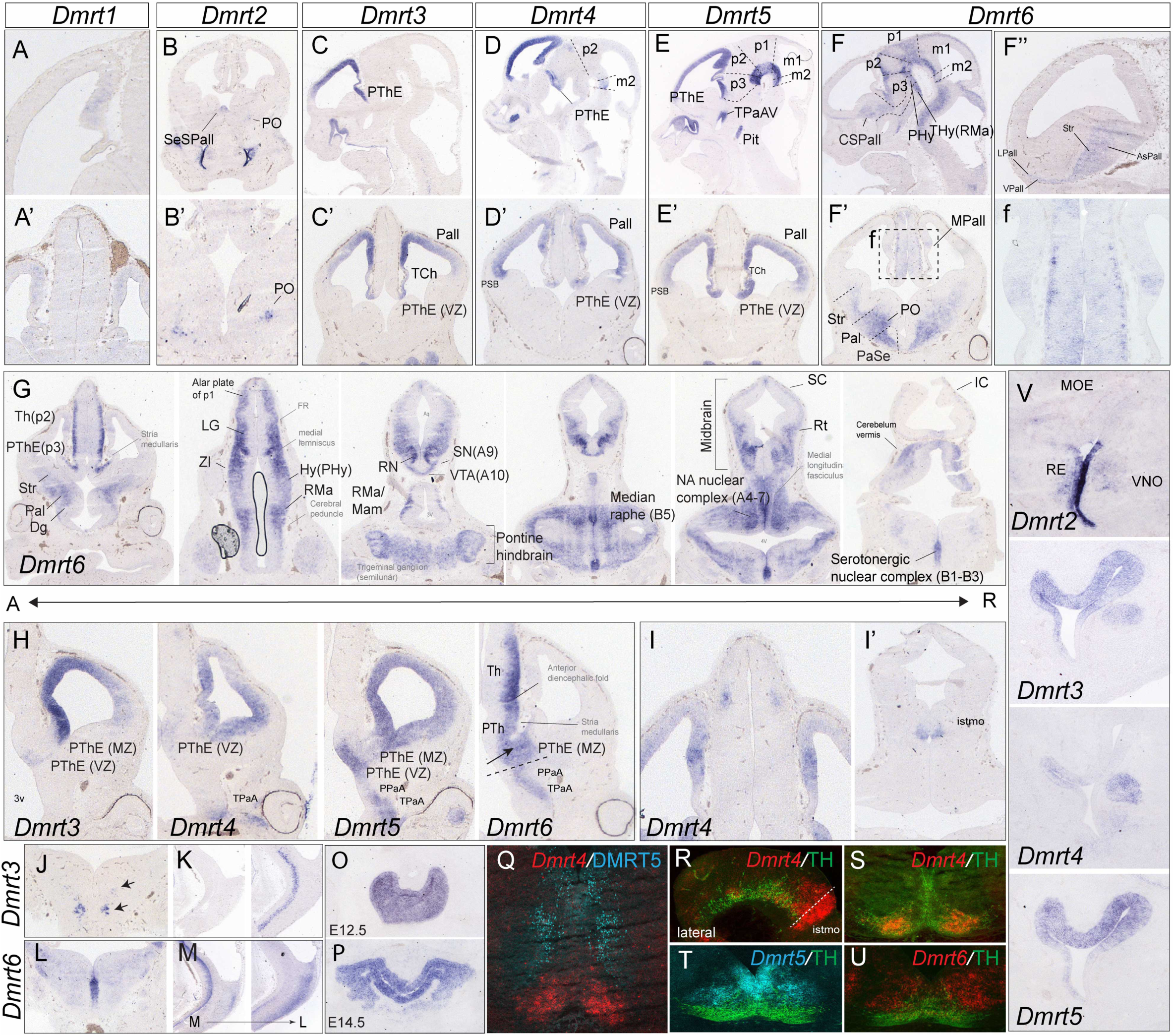
*Dmrts* are broadly expressed in early embryonic development. All panels depict ISH at E12.5 unless indicated. Only one sex is shown *Dmrt1* in the MPall in sagittal (**A**) and coronal (**A’**) views; **B-B’**) *Dmrt2* in the SeSPall and PO. **B’)** Higher magnification of PO expression. **C-E’)** *DMA-Dmrts* extensively overlap in the pallium and the PThE: *Dmrt3* (**C-C’**), *Dmrt4* (**D-D’**) and *Dmrt5* (**E-E’**). Sagittal view in upper panels, and coronal view in lower panels. **F-G**) *Dmrt6* is detected broadly in subpallial regions (CSPall, Str, Pal, PaSe) and other regions. **f)** Inset shows a magnification of the MPall region. **G**) *Dmrt6* expression along the anteroposterior axis. **H)** *Dmrt3, 4, 5* are expressed in the VZ and MZ of the PThE. *Dmrt6* is excluded from the PThE VZ. **I-I’)** *Dmrt4* is transiently expressed in the epithalamus (**I**) and the isthmus (**I’**). **J, K)** *Dmrt3* expression in the hindbrain in a coronal plane (**J**), and in a sagittal view (**K**). **L-M)** *Dmrt6* in the VZ of the B1-B3 serotonergic nuclei. *Dmrt3* and *Dmrt6* occupy distinct anatomical areas in the hindbrain, which is evident in coronal (**J, L**) and lateral views (**K, M**). **O-P**) *Dmrt5* in the adenohypophysis primordium at E12.5 (**O**) and E14.5 (**P**). **Q-U)** *Dmrt* expression in the ventral midbrain: **Q**, *Dmrt4* (ISH) and DMRT5 (IF) shows the lack of overlap between the two TFs in the ventral midbrain. **R-S)** *Dmrt4* (ISH) and TH (IF) in a sagittal view (**R**) and a coronal view (**S**). *Dmrt4* is strongly expressed in the isthmus (**R**), and it is detected in mature DA-TH+ neurons (**S**). **T and U)** *Dmrt5* and *Dmrt6* (ISH) and TH (IF). *Dmrt5* occupies the VZ, while *Dmrt6* is excluded from the VZ. None of them overlaps with TH. *Dmrt6* fills the lateral ventral domain. **V**) *Dmrt* expression in the olfactory epithelium primordium. *Dmrt2* is strongly detected in the RE and scattered cells in the VNO; *Dmrt3* and *Dmrt5* show an overlapping pattern of expression in the MOE but not in the VNO, where only *Dmrt3* is found. *Dmrt4* is strongly detected in the lateral RE and, like *Dmrt3*, it fills up the entire VNO primordium. For a complete series of expression, see: https://www.informatics.jax.org/reference/J:364871.

**Table 1.**
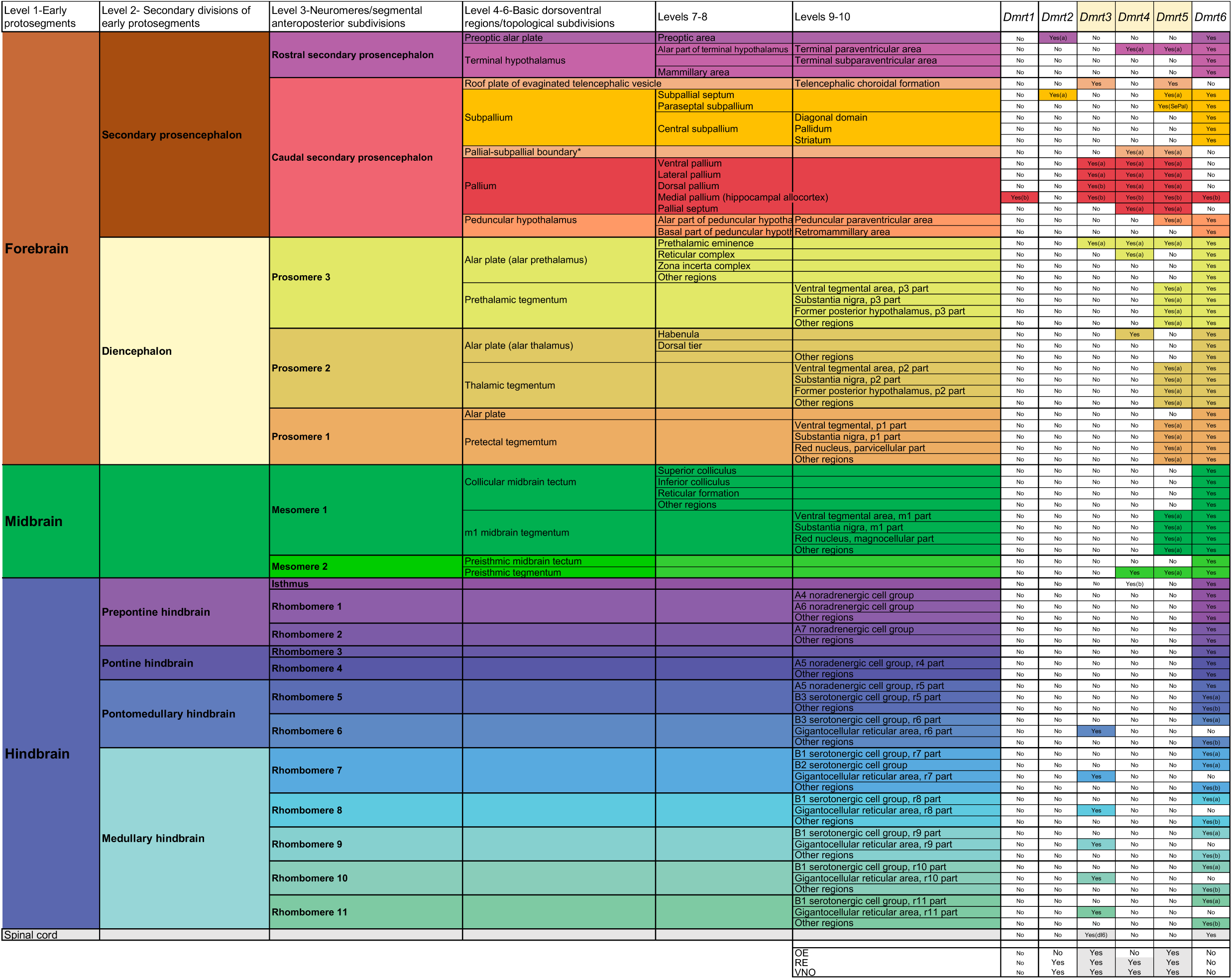
*Dmrt* expression in E12.5 embryos. (a) ventricular zone; (b) ventricular and mantle zones. Orange boxes = DMA-*Dmrts*; Grey boxes = positive expression. Presence does not necessarily mean ubiquitous expression in the nucleus.

Our comprehensive analysis revealed that *Dmrt7* was not detected in the NS (at any of the examined developmental stages). In contrast, *Dmrt2, Dmrt3, Dmrt4, Dmrt5*, and *Dmrt6* showed robust expression in the NS. *Dmrt1* was dimly detected in the medial pallium (MPall) at E12.5 (**Fig. 1A**), but no longer detected after this time point. *Dmrt2* expression was restricted to the subpallial septum (SeSPall) and the preoptic area (PO) (**Fig. 1B**). However, it was strongly expressed in the respiratory epithelium (RE) and scattered cells in the vomeronasal organ (VNO) (**Fig. 1V**). Previous reports indicate that *Dmrt6* is expressed in the mouse embryonic brain by RT-PCR^25^, but, to our knowledge, no spatial detection has been conducted in the brain before. Here, we show that *Dmrt6* is one of the most broadly expressed *Dmrts* in the mouse NS (**Table 1**), encompassing the three major parts of the brain (forebrain, midbrain, hindbrain) and the spinal cord (**Fig. 1F-1H**, **1L**, **1U**). *Dmrt6* expression was absent from proliferative ventricular zones (VZ), except in the MPall (**Fig. 1F’**) and serotonergic cell groups in the hindbrain (**Fig. 1G,1L**). This expression pattern suggested a potential postmitotic role for *Dmrt6*.

The expression of DMA-*Dmrt* members has been extensively described before, with special attention to the pallium^22^ (**Fig. 1C-E**). In addition to the three DMA-*Dmrts*, *Dmrt1* and *Dmrt6* were transiently expressed in the MPall (**Fig. 1A**, **1F’**), where the hippocampus and dentate gyrus (DG) originate from.

For the ChPl, we confirmed that *Dmrt3* and *Dmrt5* are robustly detected ^26^, and lacked *Dmrt2, Dmrt4,* and *Dmrt6* expression (**Fig. 1B, 1D, 1F**). All DMA*-Dmrts* were redundantly expressed in the VZ of the prethalamic eminence (PThE) (**Fig. 1H**), and only *Dmrt6* was found in the mantle zone (MZ) of the PThE. In addition, *Dmrt4, Dmrt5,* and *Dmrt6* were expressed in the terminal paraventricular area of the hypothalamus (TPaA). The greatest extent of *Dmrt5* in this area overlapped with a small ventral *Dmrt4* territory, and a dorsal *Dmrt6* territory (**Fig. 1H**). Moreover, *Dmrt5* and *Dmrt6* were extensively expressed in the peduncular paraventricular area of the hypothalamus (PPaA). *Dmrt5* expression was very broad and found in the ventral part (or tegmentum) of the three diencephalic (p3, p2, p1) and the two mesencephalic (m1, m2) neuromeres. This included the diencephalic portion of the former posterior hypothalamic area (diPHA), the ventral tegmental area (VTA), the substantia nigra (SN), and the red nucleus (RN) (**Fig. 1E**). *Dmrt6* was found in the same tegmentum of p3, p2, p1, m1 and m2, but, in addition, expanded to the dorsal part (or tectum) (**Fig. 1F**). *Dmrt6* continued anteriorly beyond p3 (relative to the anteroposterior axis) in both the mammillary (Mam, within the THy) and retromammilary (RMa, within the PHy) areas (**Fig. 1F**). From here, *Dmrt6* was a continuum until the hindbrain. *Dmrt5* overlapped with *Dmrt4* in the pre-isthmic tegmentum of m2 (PIsTg), where a few dopaminergic (DA) ventral mesencephalic neurons originate. In this region, *Dmrt4* was expressed in early-born postmitotic DA—as it colocalized with tyrosine hydroxylase (TH)—and *Dmrt5* lay dorsal to the *Dmrt4+* cells in the VZ and the subventricular zone (SVZ), corresponding to DA progenitors (**Fig. 1Q**–**1T**). *Dmrt6* was also expressed in the PIsTg, covering a more extensive area and partially overlapping with the domains of *Dmrt4* and *Dmrt5* (**Fig. 1U**).

We also focused on the nasal and vomeronasal epithelia due to their relevance in rodent sex-typical behaviors^27^. *Dmrt3* and *Dmrt5* overlapped almost entirely in the main olfactory epithelium (MOE) and the RE (with *Dmrt2*). In the RE, *Dmrt2* and *Dmrt4* were expressed in the ventral and lateral parts, respectively. In the VNO, *Dmrt2* presented a scattered pattern, while *Dmrt3* and *Dmrt4* filled up the whole structure (**Fig. 1V**). Previous studies have demonstrated that *Dmrt* can exhibit functional redundancy, often compensating for one another^28–30^. Therefore, exclusive sites of expression become significantly relevant in helping to identify the unique functions of each *Dmrt*. *Dmrt3* was detected in the hindbrain (**Fig. 1J**), where it has been suggested to be expressed in the dB4 neurons of the hindbrain^31,32^. However, to our knowledge, this assumption was based on the common developmental origin between dI6 and dB4 neuronal lineages and lacks a more in-depth analysis using molecular markers^32^. *Dmrt6* was also present in the hindbrain but in the VZ of serotonergic nuclei (**Fig. 1L**) and did not occupy the same territory as *Dmrt3*. Lastly, *Dmrt4* was expressed in the habenula (**Fig. 1I**) and the tegmentum of the pre-isthmus and isthmus (**Fig. 1I’**). However, similar to the other areas of the brain, *Dmrt4* was only transiently expressed in these regions. No expression was detected after E14.5 (E14.5 expression is accessible at the MGI repository, https://www.informatics.jax.org/reference/J:364871).

Although this study did not explore tissues outside the NS, *Dmrt5* expression in the developing adenohypophysis was highlighted due to its relevance to hormonal secretion, including the control of sex hormone initiation during puberty. *Dmrt5* was found from E12.5 until E18.5, as shown below (**Fig. 1O**, **1P**, **5-7, Suppl. Fig. 5**). Postnatal animals were not explored.

The analysis of the whole family in the early embryo revealed a broad representation of these genes in the mouse NS. The expression of *Dmrt1*, *Dmrt2*, and *Dmrt6* in the mouse NS was identified for the first time. At this early developmental stage, DMA-*Dmrts* were primarily found in progenitors in the VZ of distinct regions, including the pallium. Still, they also expanded to the ventral midbrain, where this early expression could be essential for its normal development.

### *Dmrt6* covers areas that control innate behaviors

*Dmrt6* has been well characterized in the male gonad, particularly in spermatogonia, and, by RT-PCR, it has been detected in the adult ovary and the NS^25,33^; however, the function of *Dmrt6* in the brain remains unknown.

From the E12.5 analysis, we concluded that *Dmrt6* was extensively expressed in subcortical structures, including the limbic system, thalamic, and brain stem structures (https://www.informatics.jax.org/reference/J:364871. The limbic system comprises distinct neural circuits of different embryonic origins that control specific innate behaviors. Sensory stimuli have a significant influence on innate behaviors, and in rodents, olfactory cues are particularly crucial, modulated by intrinsic factors such as sex. *Dmrt6* was present in the primary structures that comprise innate circuitry: the olfactory system, amygdala, bed nucleus of the stria terminalis (BST), and hypothalamus (**Fig. 2A-2J**, pink-labeled nuclei). **Table 2** complements **Figure 2**, and compiles a complete list of the regions where *Dmrt6* was maintained in late gestational embryos at E18.5 and postnatal P7 and P56 brains; **Suppl. Fig. 2**).

**Figure 2.**
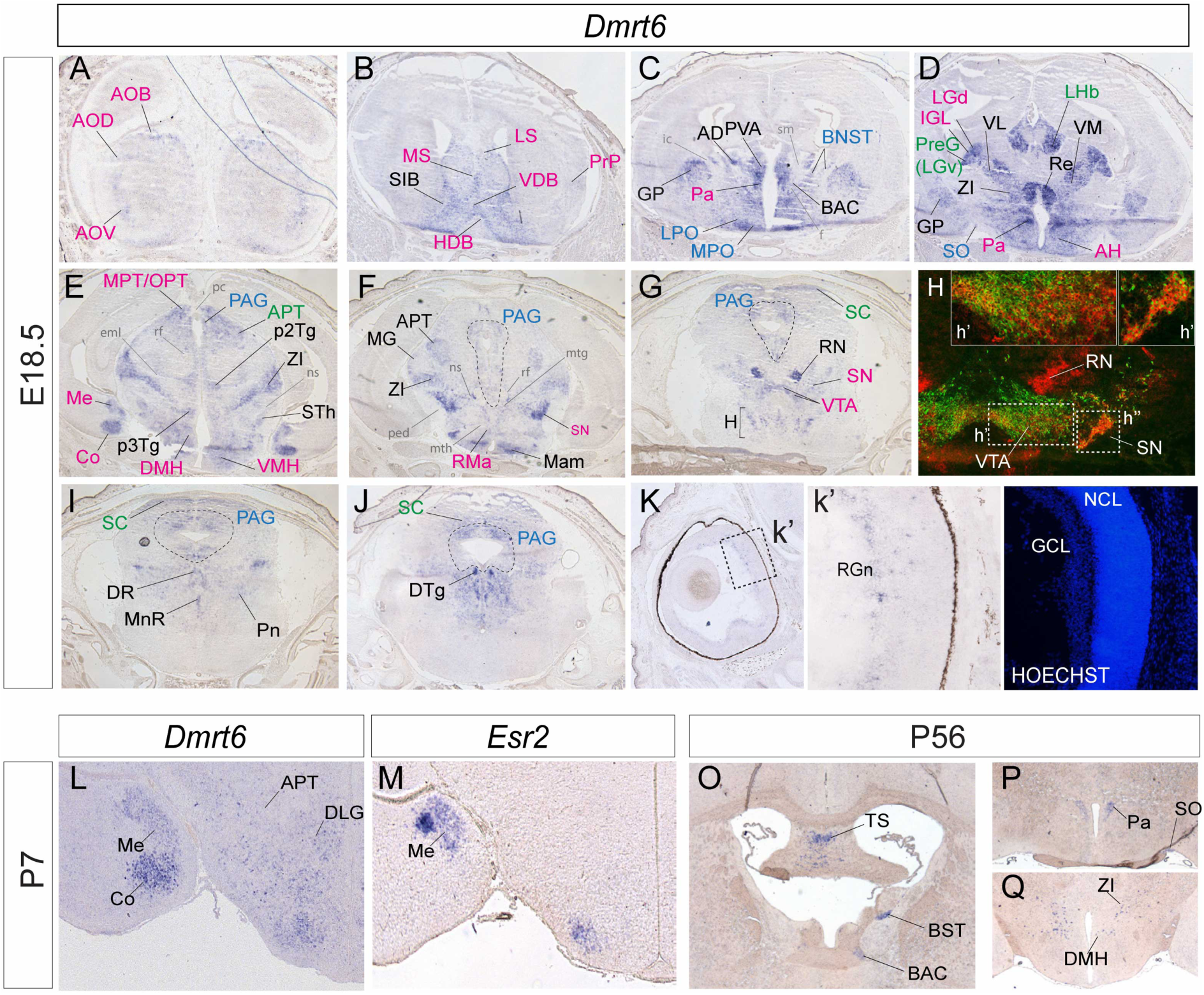
*Dmrt6* is maintained in the mouse NS, including postnatal brains. **A-K)** *Dmrt6* ISH at E18.5. Abbreviations in pink are associated with innate behaviors^34^, in green with non-image visual circuits^37^, and blue are shared. **H)** Colocalization of *Dmrt6* with TH in the VTA (**h’**) and SN (**h’’**). **K**) *Dmrt6* expression in the GCL of the retina, (**k’**) high magnification. **L)** *Dmrt6* ISH at P7 in the Me occupies the *Esr2*+ region (shown in **M**). **O-Q**) *Dmrt6* is restricted to the TS, the Pa, SO, and the DMH in adult brains.

**Table 2.**
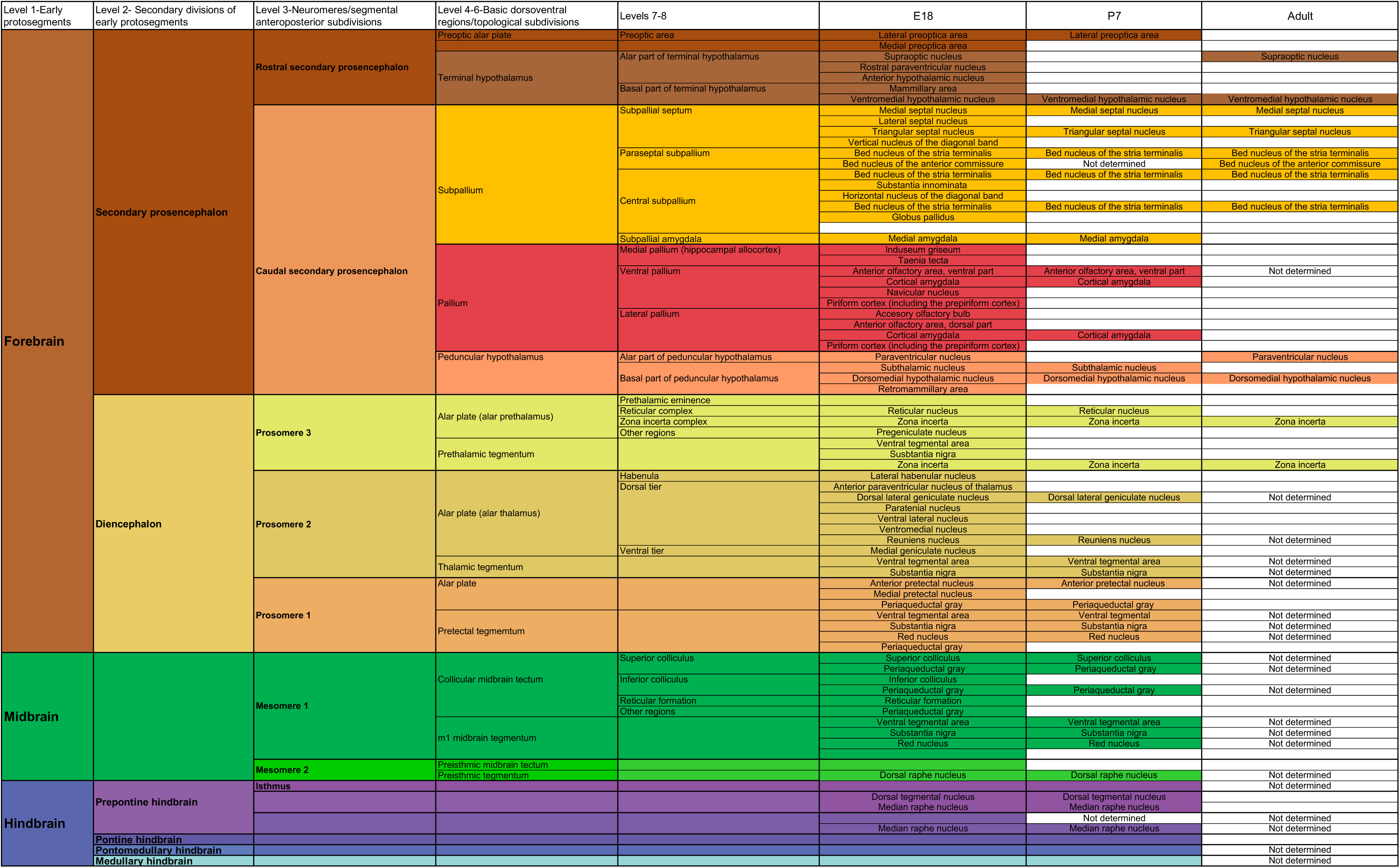
*Dmrt6* expression across development.

At E18.5, as development progresses, *Dmrt6* was detected in olfactory nuclei such as the accessory bulb (AOB), anterior olfactory nuclei (AOD, AOV), taenia tecta (TTe), and cortical amygdala (Co), one of the main limbic structures. *Dmrt6* was located in the lateral (LS) and medial (MS) septal complexes, BST, and medial amygdala (Me), which integrates many rodent sex-typical behaviors and relays information from the olfactory system to the BST^34^. Postnatally, *Dmrt6* coincided with *Esr2* (estrogen receptor beta) in the same region, suggesting a potential interaction between them (**Fig. 2L-M**). *Dmrt6* covered the PO(functionally included as part of the limbic system), and almost the whole hypothalamus, including the anterior (AH), dorsomedial (DMH), and ventromedial (VMH) hypothalamic nuclei. It overlapped with TH+ neurons in the SN and the VTA (**Fig. 2H**), where it remained postnatally animals (**Suppl. Fig. 2**). Lastly, the periaqueductal grey (PAG) is another critical region for innate behaviors where *Dmrt6* was found.

### *Dmrt6* expression in areas of the central NS related to photosensation

At E18.5, *Dmrt6* was located in the incipient ganglion cell layer (GCL) of the retina (**Fig. 2K**). From single-cell RNA sequencing data, *Dmrt6* is probably found in the intrinsically photosensitive retinal ganglion cells (ipRGCs), which are rare mammalian photoreceptors essential for a wide array of visual functions, including circadian photoentrainment, pupillary light reflex, mood, and sleep, but not for image-forming visual functions^35–37^. Several articles have described ipRGCs in non-visual circuits, as demonstrated through tracing experiments in rodents. These retinal cells connect to several brain regions through the retino-hypothalamic tract^38^, and *Dmrt6* was present in many of them: the lateral geniculate complex, including the dorsal lateral geniculate nucleus (DLG), intergeniculate leaflet (IGL), and pregeniculate nucleus (PreG) (also known as the ventral lateral geniculate nucleus); pretectal region, with the medial, olivary and anterior pretectal nuclei (involved in pupillary light reflex); lateral habenula (LHb), involved in mood regulation; BST, PO; hypothalamic nuclei, such as the supraoptic nucleus (SO); superior colliculus (SC), and PAG^37^. *Dmrt6* was excluded from the posterior and medial thalamic regions (**Fig. 2A-2J**, green-labeled nuclei), as well as from the suprachiasmatic nucleus, the central pacemaker of circadian rhythms.

*Dmrt6* was strongly expressed in other thalamic nuclei, such as the reuniens nucleus (Re), paraventricular nucleus (PVA), ventral lateral thalamic nucleus (VL), and zona incerta (ZI) (**Fig. 2A-2J**. **Table 2**). Another structure that stands out because of the strong *Dmrt6* signal is the RN (**Fig. 2G**). In the hindbrain, *Dmrt6* was maintained in the dorsal raphe (DR) and median raphe (MnR) nuclei, and pontine reticular nucleus (Pn) at E18.5 and P7 (**Fig. 2J; Suppl Fig. 2**). However, additional markers are needed to disambiguate *Dmrt6*+ cell identities in the hindbrain.

Our results, together with previous RT-PCR data^25^, confirm that Dmrt6 is broadly expressed in the NS, including regions associated with innate behaviors. Despite this, a reported *Dmrt6* conditional mouse model did not exhibit overt NS-related behavioral abnormalities^33^. The *Dmrt6* locus is highly complex with multiple transcriptional variants, some of which may lack the DM domain (NCBI gene ID: 56296; **Suppl. Fig. 1**). Thus, the allele used in Zhang et al^33^ may not have disrupted the variant expressed in the nervous system, or *Dmrt6* may influence subtle neuronal functions detectable only through specific behavioral assays, such as the pupillary light reflex test to interrogate photosensitivity.

### *Dmrt2* is maintained in the mouse NS

At E12.5, *Dmrt2* exhibited a restricted expression (**Fig. 1B**). However, its expression began to be strongly present in many regions within the cortex from E13.5 on^39^ (https://www.informatics.jax.org/reference/J:364871). **Figure 3** shows brains at P7, although the same areas were identified at E18.5 and adulthood (**Suppl. Fig. 3**; **data not shown**). According to previous studies, *Dmrt2* is expressed in corticothalamic projection neurons (CThPNs), which are specifically excluded from the somatosensory cortical area^39,40^, although no other regions have been specifically characterized. *Dmrt2* was found in the neocortex [in the orbital (ORB), prelimbic (PLC), and infralimbic (Ila) areas] and the cingulate cortex (CCx)^39^ extending into the retrosplenial cortex (RSP) (**Fig. 3A**). In addition, *Dmrt2* was found in other areas of the cerebral cortex, such as the claustrum (Cl), insular cortex (IA), and endopiriform nucleus (EPd) in a few scattered cells, and the entorhinal cortex (ENT) (also forming a continuum with perirhinal-ectorhinal cortex, PER-ECT). Lastly, *Dmrt2* was detected in the cortical amygdala (including the basomedial amygdaloid nucleus, BMA, and cortical amygdaloid area, Co). Apart from the prominent presence in cortical regions, *Dmrt2* was found in the lateral septal nucleus in its caudal part (LSc), in a few scattered cells located in the PAG (more obvious at E18.5, **Fig. 3B**), and in the pyramidal layer of the subiculum (SUB).

**Figure 3.**
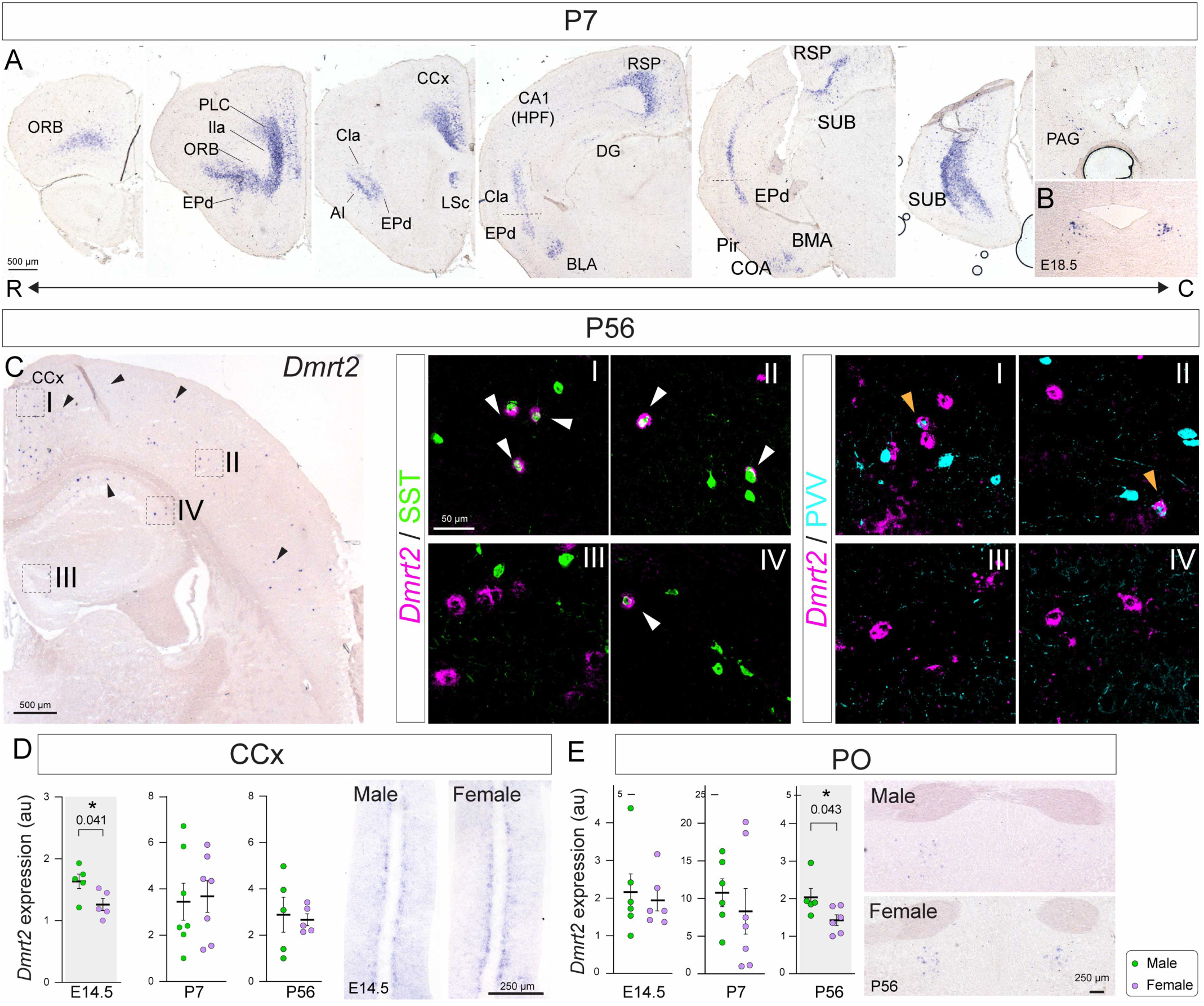
*Dmrt2* expression in postnatal brains. *Dmrt2* was detected with ISH. **A)** Expression at P7. Only one sex is shown. **B)** Expression in the PAG at E18.5 is more robust than at P7. **C)** Expression at P56. Arrowheads point to *Dmrt2*+ cells. Colocalization of *Dmrt2* with SST or PVV GABAergic markers by IF in the CCx (**I**), the neocortex (**II**), the DG, and the hippocampus (**IV**) of adult (P56) brains. White arrowheads indicate colocalization of *Dmrt2* with SST, and orange arrowheads with PVV. **D, E)** *Dmrt2* expression levels in the CCx (Adapted from^39^) and PO throughout development. The graph represents mean±SEM in arbitrary units (au). Each dot corresponds to one individual embryo. Two-tailed t-test: * p-value < 0.05 (n ≥ 5). Images show CCx at E14.5 (adapted from^39^)(**D**) and PO at P56 (**E**). R, rostral; C, caudal.

In adult brains, *Dmrt2* expression was evident in the hippocampal formation (HPF), CA1 field, and DG. This expression was more prominent in adults, although it was already detected in E18.5 and P7 (**Suppl. Fig. 3; data not shown**). Additionally, *Dmrt2* was strongly detected in somatostatin (SST) GABAergic interneurons, confirmed by *Dmrt2* ISH on adult *SSTCre;TdT* brains (**Fig. 3C**). Most *Dmrt2*+ cells colocalized with SST, especially in the medial part of the neocortex, where almost every *Dmrt2*+ neuron was SST+. We found less colocalization between SST and *Dmrt2* in the lateral cortex (**Fig. 3Ci-iv**), and no colocalization with PVV anywhere, except for some anecdotal neurons. A similar situation was found for *Dmrt2* neurons in CA1, where a few colocalize with SST and none with PVV. *Dmrt2* did not colocalize with any of these markers in the DG. This finding aligns with the Allen Brain Single Cell Database and https://scheiffele-splice.scicore.unibas.ch/.

### *Dmrt2* expression levels differ between the sexes in the CCx and PO

We have previously shown that *Dmrt2* expression levels in the CCx are higher in males than females during embryonic development^39^. In contrast, this differential expression is lost in postnatal and adult brains (**Fig. 3D**). The expression of *Dmrt2* in the PO of early embryos (**Fig. 1B**) and adult (**Fig. 3E**) prompted us to focus on this region. The PO is one of the areas in rodents associated with innate functions, such as sexual and parental behaviors, and presents molecular sex differences in cell numbers and dendritic density, which are controlled by gonadal hormones^41^. We dissected the POs of male and female embryos at E14.5, P7, and P56, and conducted RT-qPCR. *Dmrt2* exhibited higher expression levels in adult males, but not at other developmental stages (**Fig. 3E**). We did not conduct cellular quantifications or identification. With our analysis, we cannot conclude whether there was a difference in *Dmrt2*+ cell numbers or higher levels of expression per cell.

Our findings indicate that distinct mechanisms may regulate *Dmrt2* expression levels. In the CCx, *Dmrt2* levels vary between sexes when hormonal secretion is low^42^. In contrast, in the PO—a region under strong regulation by sex hormones—*Dmrt2* expression increases in males following puberty, coinciding with the establishment of hormonal secretion in sexually mature animals. Further research is required to elucidate the origins of these differential *Dmrt2* expression patterns and their physiological significance. Unfortunately, the study of *Dmrt2* loss-of-function mutants is limited to embryonic stages due to their perinatal lethality^43^.

### Persistent expression of DMA-*Dmrts* beyond early embryonic stages

DMA-*Dmrts* expression in the NS has been described in detail at early embryonic stages^19,22,26,44–46^ (**Table 1**, **Fig. 1C-2E**, https://www.informatics.jax.org/reference/J:364871). To our knowledge, whether their expression persists in any brain region has not been documented before. We found that among the DMA-*Dmrts*, only *Dmrt3* and *Dmrt5* were maintained, while *Dmrt4* was no longer detected anywhere in the mouse brain after E14.5 (https://www.informatics.jax.org/reference/J:364871).

We highlighted the expression of DMA-*Dmrts* in non-neural structures, which are closely related to brain function. In the ChPl, *Dmrt3* and *Dmrt5* were strongly detected across development from early embryos until adults (**Fig. 1C’-1E’**, **Fig. 4K**, **4L**). Both TFs were localized to epithelial cells of the lateral ventricle ChPl, but not in those of the third or fourth ventricles, which have different developmental origins. It has been previously described that *Dmrt5* is necessary for the formation of the ChPl, although this might be a consequence of *Dmrt5* expression in the cortical hem^26,44,47,48^.

**Figure 4.**
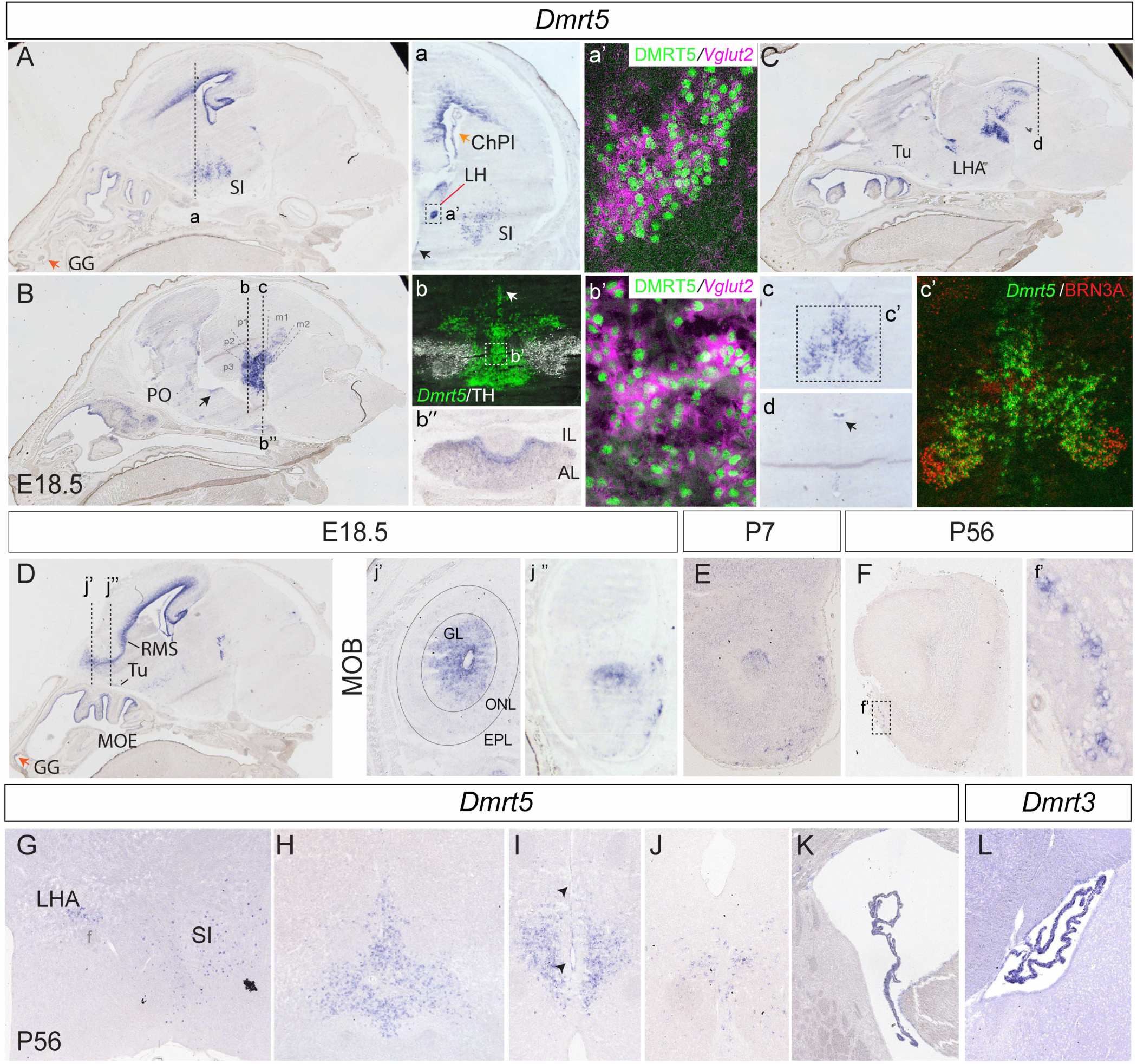
*Dmrt5* is maintained in postnatal brains. *Dmrt5* was detected with ISH, except in **a’** and **b’**, where double DMRT5 IF and *Vglut2* ISH was performed. **A, B, C,** and **D)** Sagittal views of *Dmrt5* at E18.5. **a**-**d** coronal views corresponding to levels indicated in **A), B),** and **C)**. Orange arrow in **A)** and **D)** indicates the location of *Dmrt5* in the GG. Black arrows in **A)**, **B**), and **I)** in the CVOs. **a’** and **b’**) DMRT5 in the LH and the VTA in glutamatergic (*VGluT2*+) neurons. **b**) *Dmrt5* does not colocalize with TH in the VTA. **c’)** *Dmrt5* colocalize with BRN3A in the RN. **D**) At E18.5, *Dmrt5* fills up the whole RMS and other olfactory system regions, the Tu, the MOE, and the GG, including the MOB (**j’** and **j’’**). It perdures in postnatal OBs (**E, F**). In adults, it is restricted to a bead-like pattern that resembles the necklace glomeruli (**f’**). *Dmrt5* expression is maintained in other regions in adulthood such as the LH and SI (**G**), the VTA (**H**), and PHA (**I**), including in the ventricle ependymal layer, black arrows (**I**). *Dmrt5* and *Dmrt3* persist in the ChPl (**K** and **L**).

*Dmrt5*, but not *Dmrt3*, was also found in the adenohypophysis, presumably expressed in progenitor cells at E12.5 and E14.5 (**Fig. 1O**, **1P**). At E18.5, *Dmrt5* was detected in the marginal cell layer (MCL) of the intermediate lobe (IL), and in a few cells of the anterior lobe (AL) (**Fig. 4Bb’’**). The role of *Dmrt5* has been described in the zebrafish hypophysis, where *Dmrt5* is required for the differentiation of corticotropes and gonadotropes, as well as for the maintenance of lactotropes, ultimately affecting reproductive function^49^. Whether this function is conserved in mammals is unknown. In line with the phenotype in zebrafish, we did not observe an overt morphological pituitary phenotype in *Dmrt5^-/-^* mice (**data not shown**).

Among the three DMA-*Dmrts*, *Dmrt5* exhibited the broadest expression pattern in early embryos (**Table 1**). It was maintained in many more nuclei in late embryonic and postnatal brains than *Dmrt3* (**Fig. 4-7**). *Dmrt5* presented a scattered expression pattern in the PO and subpallial structures, such as the olfactory tuberculum (Tu) (**Fig. 4C**,**4G**), the vertical nucleus of the diagonal band (VDB) (**Suppl. Fig. 4**), and the SI (**Fig. 4A**, **4G**). Furthermore, *Dmrt5* was expressed in the island of Calleja major (ICjM), a postnatally born structure (**Suppl. Fig. 4**), and the lateral hypothalamic area (LHA) (**Fig. 4a’**,**4C**).

*Dmrt5* encompassed a plurisegmental region from the diencephalon to the midbrain tegmentum (**Fig. 1E**)^44,50^. In early embryos, *Dmrt5* was expressed in midbrain LMX1A+ DA progenitors and extended laterally to the glutamatergic m6 domain (**Fig. 1T**)^50^. The continuous *Dmrt5*+ domain is largely glutamatergic (colocalizes with *Vglut2*) and persists through development until adulthood along the anteroposterior axis covering the diPHA (**Fig. 4Bb, b’**, **4H**, **4I**) and several postmitotic mesencephalic nuclei: the VTA, with minimal overlap with DA (TH+) cells (**Fig. 4Bb**); the RN, overlapping with BRN3A+ (POU4F1) cells (**Fig. 4Bc’**); the Edinger-Westphal nucleus (EW); the midbrain reticular nucleus (MRN), and the PAG (**Fig. 4Bd**).

In the ventral midbrain, *Dmrt5* plays an early role in patterning, in addition to regulating DA identity at the expense of GABAergic neurons. *Dmrt5* suppresses the transcriptional program controlling the ventral-lateral midbrain and promotes ventral-medial progenitor fate (*Msx1, Lmx1a, Foxa2*)^50^. It would be interesting to test whether *Dmrt5* regulates DA neurons *in vivo* and their postmitotic function.

### *Dmrt5* is enriched in olfactory structures across development

Apart from MOE and VNO, *Dmrt5* was maintained in other olfactory structures, such as the Grueneberg ganglion (GG) (**Fig. 4A**, **4D**), the main olfactory bulb (MOB) (**Fig. 4Dj’**, **4Dj’’**), and the Tu (**Fig. 4C**, **4D**). *Dmrt5* appeared in the MOB glomerular layer (GL) in the late embryo (**Fig. 4Dj’**). At P7, MOB layers are fully formed, and *Dmrt5* expression was maintained in the GL. However, the expression is more restricted than at E18.5 (**Fig. 4E**). In the adult, expression of *Dmrt5* in the GL is limited to a few glomeruli of the ventrolateral zone of the MOB (**Fig. 4Ff’**). The bead-like expression pattern resembles the necklace glomeruli. However, specific markers are needed to confirm this type of glomeruli, such as *Gucy2d* or *Pde2a*^51^. The embryonic function of *Dmrt5* in the olfactory system will be approached later in this manuscript.

### Dmrt3 and Dmrt5 persist in postnatal neurogenic niches

By analyzing late embryos and postnatal animals, we observed that *Dmrt5*, and *Dmrt3* to a lesser degree, persisted in postnatal neurogenic niches, including the SVZ, the DG, and the MOE. Furthermore, *Dmrt5* expression appeared to be detected in circumventricular organs (CVOs), which comprise a midline series of adult stem cell niches along the third and fourth ventricles^52–54^, in the ventral side of the third ventricle (potentially the vascular organ of the lamina terminalis, VOLT) (**Fig. 4Aa**, **4B**, black arrows), where it persisted in adults (**Fig. 4I**, black arrows), and the area postrema (AP) (**Fig. 4C**, **4Bd**).

At E12.5, *Dmrt3* and *Dmrt5* are prominently expressed in neocortical progenitor cells in the pallial VZ (**Fig. 1C,1E**)^22^. At E18.5, those progenitors are depleted; therefore, DMA-*Dmrts* expression was not detected in the cortex. However, the SVZ persists after VZ depletion^55^. Concomitantly, *Dmrt5* expression remained in the dorsal and septal SVZ (not the striatal) in its entire anteroposterior extension (**Fig. 4A**, **4D**), following the rostral migratory stream (RMS) (**Fig. 4D**). In the OB, *Dmrt5* filled the olfactory ventricle and the SVZ. We could still detect *Dmrt5* mRNA (but not the protein) in adult animals in the dorsal SVZ (**Fig. 5A**, dashed line demarked by black arrowheads).

**Figure 5.**
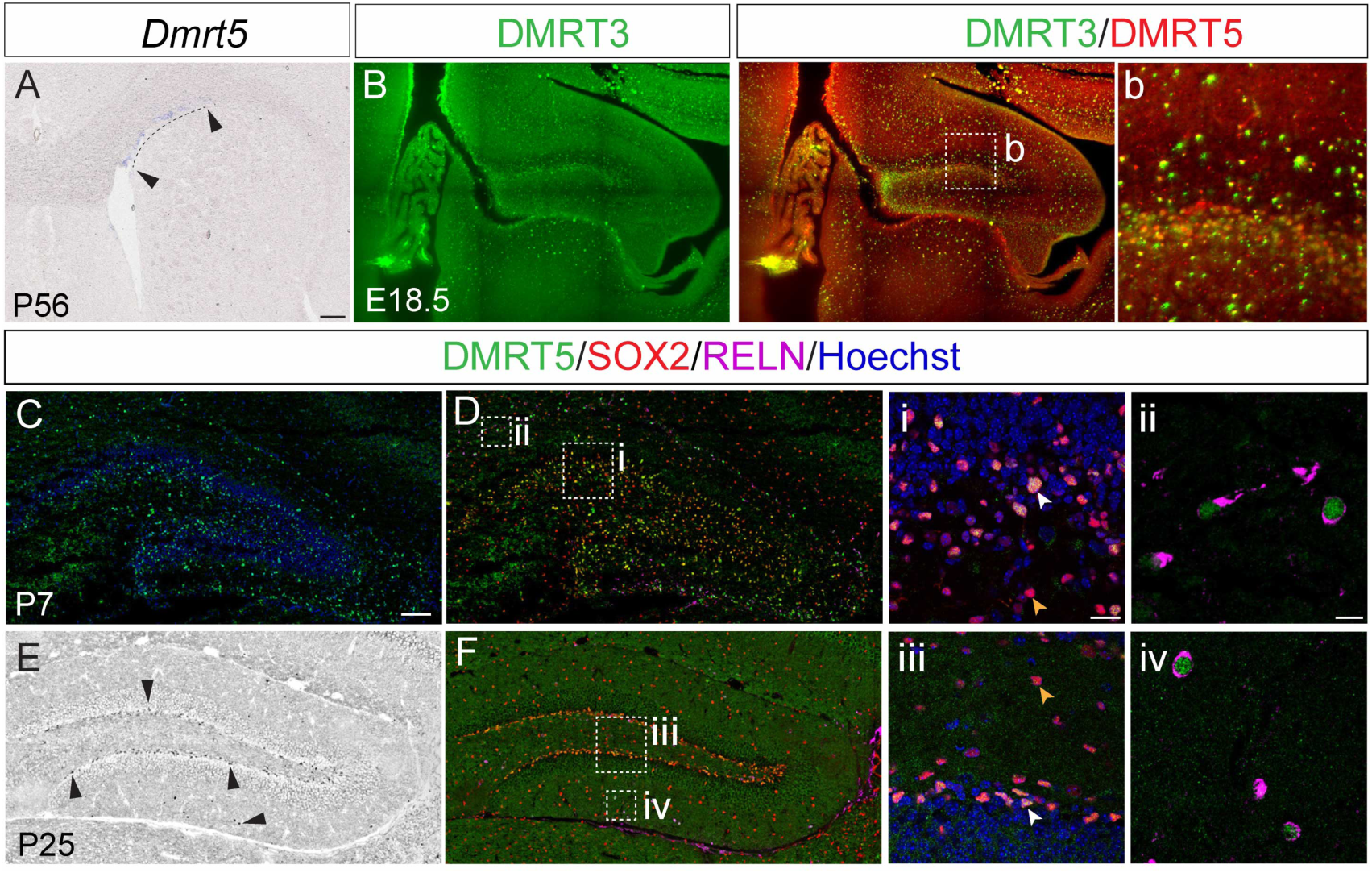
DMRT3 and DMRT5 in the postnatal neurogenic niches. **A)** *Dmrt5* mRNA in the dorsal SVZ in adults (dashed line demarks the *Dmrt5* area). **B)** DMRT3 and DMRT5 overlap in the DG—coronal section from a Light sheet scan obtained from an iDISCO-treated E18.5 brain. **b**) High magnification of the SGZ. Bright yellow dots are an artefact of the treatment. **C, D)** At P7, DMRT5 (IF) colocalizes with SOX2 in the hilus (**i**) and REELIN in CR cells (**ii**). In **C)** DMRT5 with nuclear Hoechst staining shows the distribution of DMRT5 in the hippocampal formation. **E)** At P25, the inverted image shows that DMRT5 (cells shown in black, marked by arrowhead) persisted in the SGZ, overlapping with SOX2 in NPCs (**F**). Insets **i)** and **iii)** show a high magnification of the SGZ. The white arrowheads depict DMRT5+/SOX2+ colocalization. The orange arrowheads show SOX2+ outside the SGZ (potentially astrocytes) that do not express DMRT5. In **ii** and **iv),** a few CR, expressing both REELIN and DMRT5, remain postnatally.

A second prominent site of postnatal neurogenesis is the subgranular zone (SGZ) of the DG, where progenitor cells remain until adulthood in the mouse brain^56^. DMRT3 and DMRT5 proteins overlapped in the E18.5 DG (**Fig. 5B**). At P7, DMRT5 colocalized with SOX2, a marker for neural stem cells and progenitor cells in the hilus^57^, corresponding to the tertiary dentate matrix (**Fig. 5C, 5D**). At P25, SOX2+/DMRT5+ cells are absent from the hilus and confined in the SGZ (**Fig. 5D**, **5E**, black arrowheads). In addition, at P7 and P25, DMRT5 colocalized in the DG with reelin (RELN), a marker for Cajal-Retzius cells (CR) (**Fig. 5ii**, **5iv**). *Dmrt5* is crucial for the generation of CR in the medial pallium^44^. In *Dmrt5^-/-^*mutants, there is a complete absence of cortical hem-derived CRs (the ones that eventually populate the postnatal hippocampus) and those from the pallia-subpallial boundary (PSB)^44,46,26^. We observed a decline in *Dmrt5* expression from P7 to P25 in the SGZ, which might correlate with the reported reduction of postnatal neural progenitors over time^58^ (**Fig. 5C**, **5E**, respectively). Another neural structure that can regenerate throughout life is the MOE^59,60^. *Dmrt3* and *Dmrt5* were maintained there postnatally, showing a strong signal in the basal layer, where olfactory neural stem cells (oNSCs) are lifelong maintained, and in the sustentacular cells of the apical layer^61^ (**Fig. 6A**, **6B**). In addition, *Dmrt3* was found in the apical layer of the sensory epithelium of the VNO (sVNO), the marginal zone, and a few cells of the basal layer of the sVNO (**Fig. 6Aa’’**). The marginal zone is located at the outer edges of the sVNO, where it interfaces with the non-sensory epithelium of the VNO (nsVNO), constituting the lifelong stem cell niche of the VNO^62,63^.

**Figure 6.**
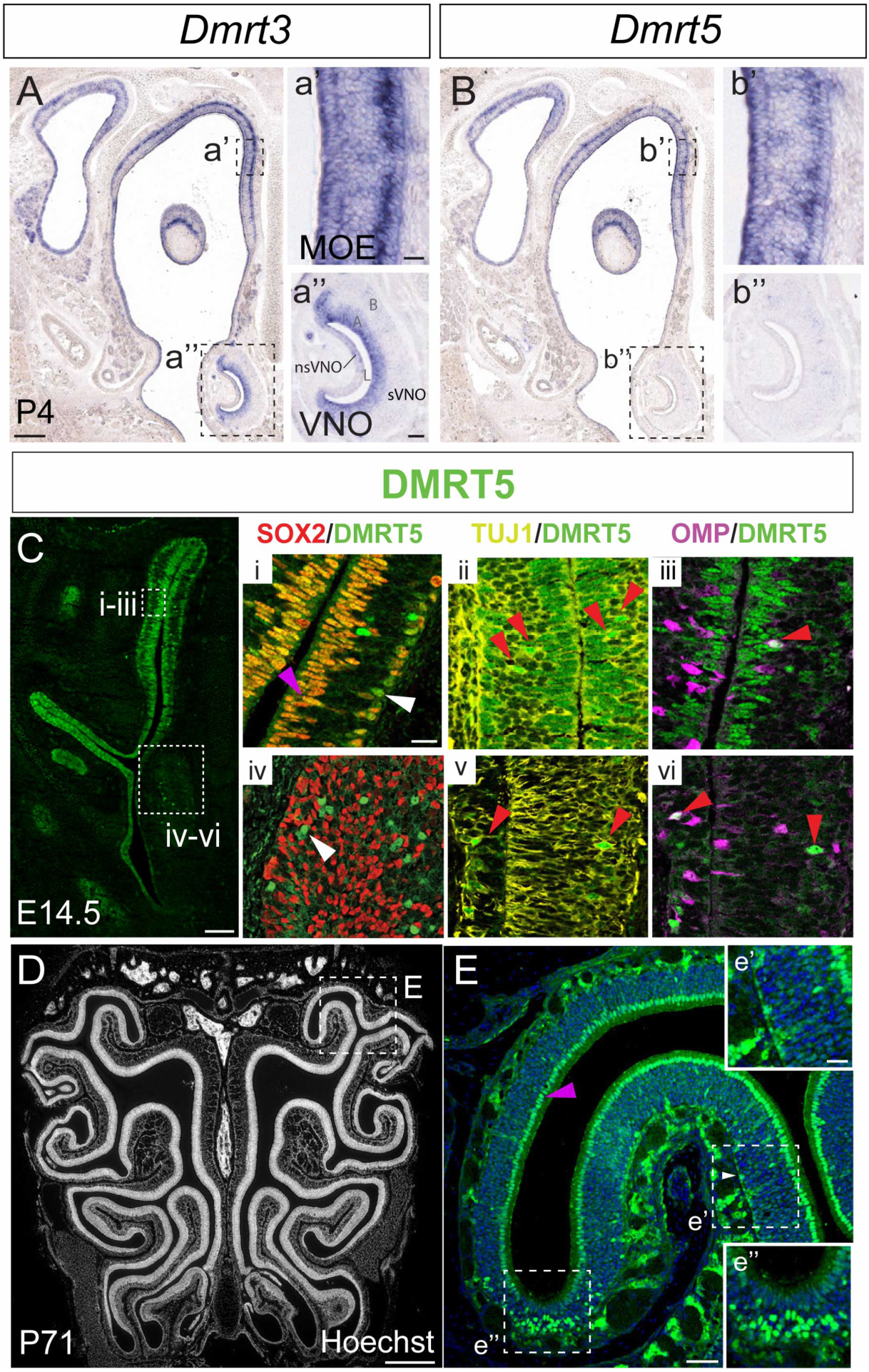
*Dmrt3* and Dmrt5 in the MOE across development. **A, B)** *Dmrt3* and *Dmrt5* (ISH) are expressed in the MOE and VNO of P4 animals. Scale bars: 200 µm (A), 20 µm (a) and 10 µm (a’). **C)** Identification of DMRT5 cellular subtypes in the MOE (**i, ii,** and **iii**) and the VNO (**iv, v,** and **vi**) at E14.5. In the MOE, DMRT5 colocalized with SOX2 in the apical layer (magenta arrowhead) but to a lesser extent in the basal layer (white arrowhead indicates a DMRT5+/SOX2-cell) (**i**). In the intermediate zone, DMRT5 colocalized with TUJ1 (immature) and OMP (mature) sensory neurons (red arrowheads in **ii** and **iii**). In the VNO, DMRT5 does not overlap with SOX2, although it does with TUJ1 (**v**) and OMP in the nsVNO (**vi**). Scale bars: 100 µm (C), 20 µm (i), 500 µm (D), 50 µm (E), and 20 µm (e’). **D, E)** DMRT5 is maintained in adult MOE. The Hoechst staining in **D)** depicts the turbinates in an adult animal. **E)** Inset of one turbinate with DMRT5 staining robustly located in the sustentacular layer (magenta arrowhead). DMRT5 remains in a few cells in the basal layer where NPCs are located (**e’**), and in the “cul-de-sac”, there is a prominent concentration of DMRT5 cells (**e’’**). A: apical layer of sVNO; B: basal layer of sVNO; L: lumen.

We characterized DMRT5+ cell types in the MOE through immunostaining (**Fig. 6C**, **6D**). DMRT5 colocalized with SOX2+ oNSCs, with a few DMRT5+/SOX2-cells likely corresponding to differentiating intermediate progenitors (**Fig. 6Ci**, white arrow). DMRT5 and SOX2 co-localization was also observed in the sustentacular cells (**Fig. 6Ci**, magenta arrow). Additionally, scattered DMRT5+ cells in the intermediate layer corresponded to immature (TUJ1+/OMP-) and mature (OMP+) olfactory sensory neurons (iOSNs and mOSNs, respectively; **Fig. 6Cii**, **6Ciii**, red arrows). This expression pattern persisted into adulthood (**Fig. 6D**), where DMRT5 continued to mark the sustentacular cells (**Fig. 6E**, indicated by the magenta arrow). In addition, a few scattered cells lay within the basal layer (**Fig. 6Ee’**, white arrow). DMRT5 was also found in the curved portions of the regions called “cul-de-sacs” (**Fig. 6Ee’’**), which are blind-ended recesses within the posterior end of the nasal cavity that contain non-canonical OSNs defined by the expression of the transmembrane receptor guanylate cyclase D and often exhibit a low level of OMP^64^. Their axons project to the necklace glomeruli of the olfactory bulb^65,66^, where *Dmrt5* has also been detected in this study (**Fig. 4g’**). They detect social cues, such as the peptide uroguanylin and the gas carbon disulfide, to mediate food-related social learning^67–69^.

Similar to the MOE, the embryonic VNO exhibited scattered DMRT5+ immature and mature olfactory sensory neurons in the sensory epithelium (**Fig. 6Bb’’**, **6Civ-vi**). However, unlike the MOE, DMRT5 was absent from the sustentacular cells of the VNO. The non-sensory VNO epithelium contains DMRT5+ cells that colocalized with TUJ1 and OMP (**Fig. 6Cv and 6Cvi**, respectively), though their function remains undetermined.

We showed that DMRT3 and DMRT5 proteins were maintained in neural progenitor cells (NPCs) in postnatally. DMRT3 was only detected in this study in the DG (no later stages were analyzed), colocalizing with DMRT5. In addition, DMRT5 was located in several postnatal neurogenic sites: the basal progenitors of the MOE, the SVZ and SGZ of the DG, and in CVOs. In all cases, DMRT5 colocalized with SOX2, becoming a novel marker of NPCs^57^.

Previous studies have shown that *Dmrt5* sustains progenitor proliferation by regulating *Hes1* and cell cycle genes, and its loss strongly impairs hippocampal and cortical development^45,70^. Similar phenotypes have also been observed in zebrafish, indicating a strong conservation across different species^71,72^. Double-knockout mice for *Dmrt3* and *Dmrt5* exhibit a more severe phenotyp^28^. If similar mechanisms were to be employed in postnatal NPCs, *Dmrt3* and *Dmrt5* might be maintaining postnatal progenitors in a quiescent state. It would be interesting to investigate in future experiments how they contribute to adult neurogenesis and whether they might integrate sexual cues, such *as Dmrt* in *Drosophila* and *C. elegans*^73^.

### *Dmrt5* is required for the development of several components of the olfactory system

The olfactory system plays a crucial role in regulating rodent innate behaviors, which often exhibit pronounced sex differences^74^. Given the robust expression of *Dmrt5* in multiple olfactory nuclei across various levels of information processing, we were prompted to investigate its function in specifying the olfactory system in both males and females at distinct circuit levels.

The olfactory system is the only sensory system in which the information reaches the primary sensory cortex without passing through the thalamus. Sensory neurons in the MOE and VNO detect pheromones and other social cues. Then, these neurons pass the information on to OB neurons that project to the olfactory cortex, which includes the Pir and the Tu. Previous data in *Xenopus laevis* indicate that *Dmrt5* is necessary for neurogenesis in the nasal placode^75^, but this has never been assessed in mammals. For our analysis, we used *Dmrt5* null mutants kindly provided by Prof. Bellefroid (*Dmrt5*^-/-^) (**Suppl. Fig. 1**)^44^. The perinatal lethality of *Dmrt5^-/-^* homozygous mice limited our study to embryonic stages.

### *Dmrt5* is required for the general olfactory epithelium specification

DMA*-Dmrts* could compensate for each other functionally^28,75^. DMRT3 and DMRT5 have similar DNA-binding properties, and in cortical development, they have a synergic effect on their target genes^28,30^. Therefore, we first established the epistatic relationship between the three *DMA-Dmrts* in the nasal epithelium by analyzing *Dmrt3* and *Dmrt4* in *Dmrt5*^-/-^ mutants. At E14.5, *Dmrt3 and Dmrt5* overlapped completely in the OE (including the MOE and the RE), while *Dmrt4* was only dimly detected in the lateral part of the RE. *Dmrt3* and *Dmrt4* showed a similar expression pattern in the VNO: filling up the entire apical layer and the marginal zone. In contrast, their expression was more restricted in the basal layer, being localized to the most basal cells (**Fig. 1V**, **8A-8C**). In *Dmrt5*^-/-^ mutants, *Dmrt4* expression was ectopically expressed in the MOE and upregulated in the RE (**Fig. 8C’**), while *Dmrt3* showed an expression similar to that of the wildtype (WT) (**Fig. 8B’**). No gross defects were found in the VNO for *Dmrt3* or *Dmrt4*. These results suggest that *Dmrt5* is repressing *Dmrt4* expression in WT conditions, following a similar logic to that of the pallium^30^.

In *Dmrt5^-/-^*, the MOE area was reduced, likely due to a decrease in the total number of cells (**Fig. 7F**, **7G**). In WT, SOX2 cells were arranged in basal and apical layers (in neural progenitors and sustentacular cells, respectively). In *Dmrt5*^-/-^ mutants, SOX2+ cells were randomly distributed across the epithelium, indicating that the MOE lost its organized layer stratification. SOX2 protein distribution was not abolished; however, the apical layer presented fewer SOX2+ cells (**Fig. 7D’**). Furthermore, in *Dmrt5*^-/-^, the neuronal markers TUJ1+ (for both iOSNs and mOSNs) and OMP+ (specific for mOSNs) were ectopically located in the RE, in its lateral and ventral parts (**Fig. 7E’b**, **7F’**). Next, we measured the mOSN density in both sexes (number of mOSN divided by the MOE area depicted in dashed red lines in **Fig. 7F**, **7F’**). Both the total number of mOSNs and the MOE area were reduced in mutants of both sexes. As a result, the mOSN density was not different between genotypes or sexes, indicating that *Dmrt5* affected the overall MOE specification similarly in males and females, and did not specifically regulate sensory neurons in the MOE (**Fig. 7G**).

**Figure 7.**
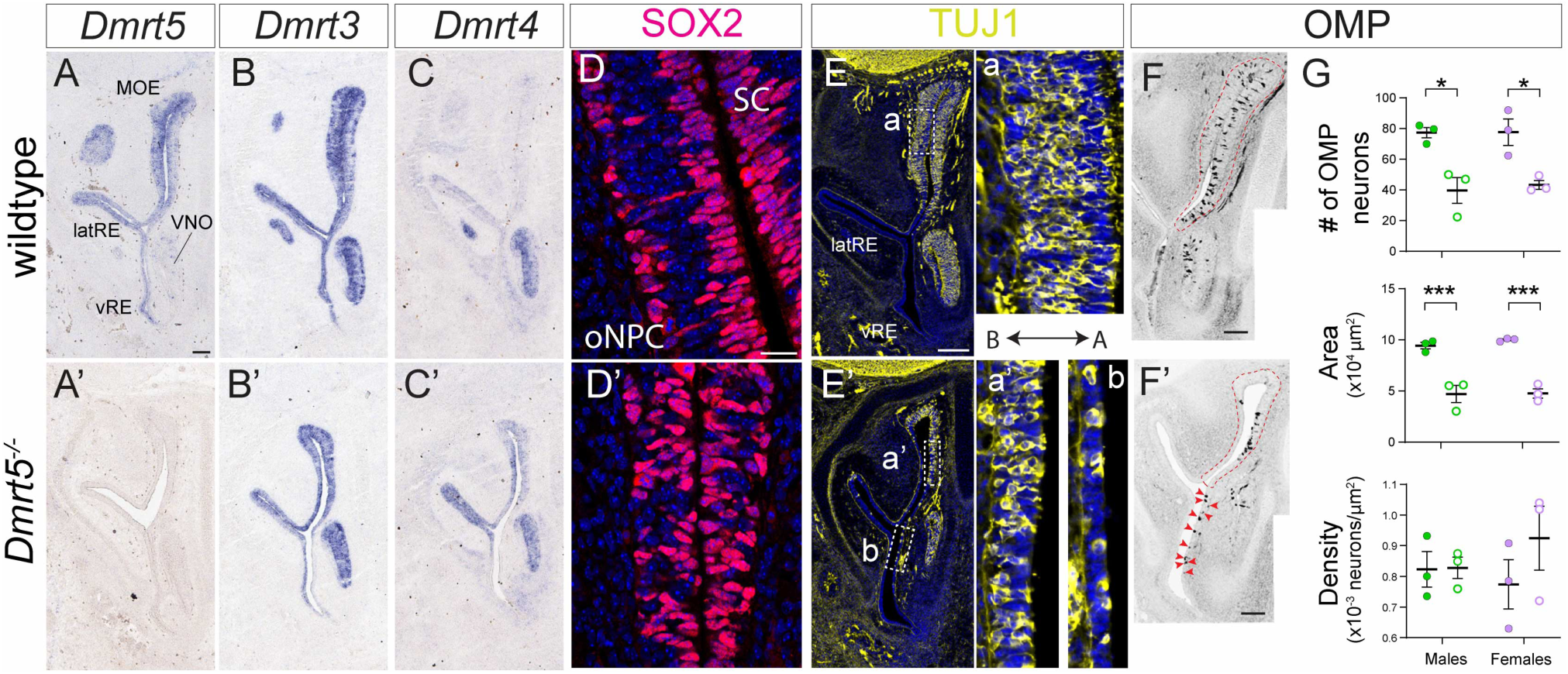
*Dmrt5* is required for the specification of the MOE in both sexes. **A, B,** and **C)** *Dmrt5*, *Dmrt3,* and *Dmrt4* ISH in WT nasal epithelium at E14.5. **A)** *Dmrt5* is not detected in *Dmrt5^-/-^*; however, the MOE is reduced. **B)** In *Dmrt5^-/-^*, *Dmrt3* ISH does not show overt differences in expression, while *Dmrt4* is ectopically expressed in the RE and upregulated in the MOE (**C**). Scale bar: 100 µm (A). **D, D’)** SOX2 staining in basal and apical layers of the MOE in WT (**D**) and *Dmrt5^-/-^* (**D’**). SOX2 cells are disorganized in the mutants. Scale bar: 20 µm. **E, E’**) TUJ1 expression in WT (**E**) and *Dmrt5^-/-^* (**E’**). TUJ1 fills the entire MOE (**a**) and the VNO in WT conditions. In *Dmrt5^-/-^*, the MOE is thinner (**a’**), although TUJ1 is maintained, and there is ectopic distribution in the RE (**b**). Scale bar: 200 µm (E) and 20 µm (a). **F, F’)** OMP distribution is shown in inverted images. There is a significant reduction in the number of OMP neurons (quantification in **G**). In addition, there are ectopic OMP+ neurons in the RE (red arrowheads). Scale bar: 100 µm. In **G)** graphs represent the mean±SEM of the number of OMP neurons in one section of the unilateral MOE, area of the unilateral MOE (dashed red line), and the OMP+ cell density. Each dot corresponds to a unilateral MOE from an individual embryo. Two-way ANOVA followed by Tukey multiple comparisons test: * p-value < 0.05; *** p-value < 0.001. B, basal; A, apical.

These results suggest that *Dmrt5* is required for the mouse MOE specification, with a similar effect in both sexes. As a result, neuronal and non-neuronal (such as sustentacular) cell types are reduced in *Dmrt5*^-/-^. Previous studies in *Xenopus* have shown that *XDmrt5* and *XDmrt4* may function redundantly upstream of the pro-neural genes *XNgnr1* and *XEbf2* in olfactory placode neurogenesis^75^. The ectopic expression of *Dmrt4* and the continued presence of *Dmrt3* in the MOE may together compensate for the absence of *Dmrt5*. To further elucidate the role of *Dmrt5*, future studies employing specific molecular markers and epistasis analyses with established TFs involved in MOE-specific processes are essential.

### Sex and *Dmrt5* interact to regulate the number of DA neurons in the OB and Pir

In early embryos, *Dmrt5* (together with *Dmrt3* and *Dmrt4*) determines the position and size of the PSB^45^ from where OB interneurons (OB-I, TH+) and OB projection neurons (OB-p, Calretinin+) derive. In fact, *Dmrt5*^-/-^ embryos show an expansion of the OB-i population^26^. Accordingly, we observed an increase in TH+ neurons in *Dmrt5*^-/-^ E18.5 embryos (**Fig. 8A-8B**). To quantify the density of TH+ neurons and investigate the potential interaction between *Dmrt5* and sex, we performed 3D brain reconstructions in clarified brains (by iDISCO+, see **Experimental Procedures**) stained with an anti-TH antibody. This technique allowed us to precisely measure the number of TH interneurons normalized by the OB volume (**Fig. 8C-8F**). The TH density was similar between the sexes in WT OBs. However, in the *Dmrt5^-/-^*background, the TH cellular density increased significantly in female mutants but not in males (**Fig. 8E**).

**Figure 8.**
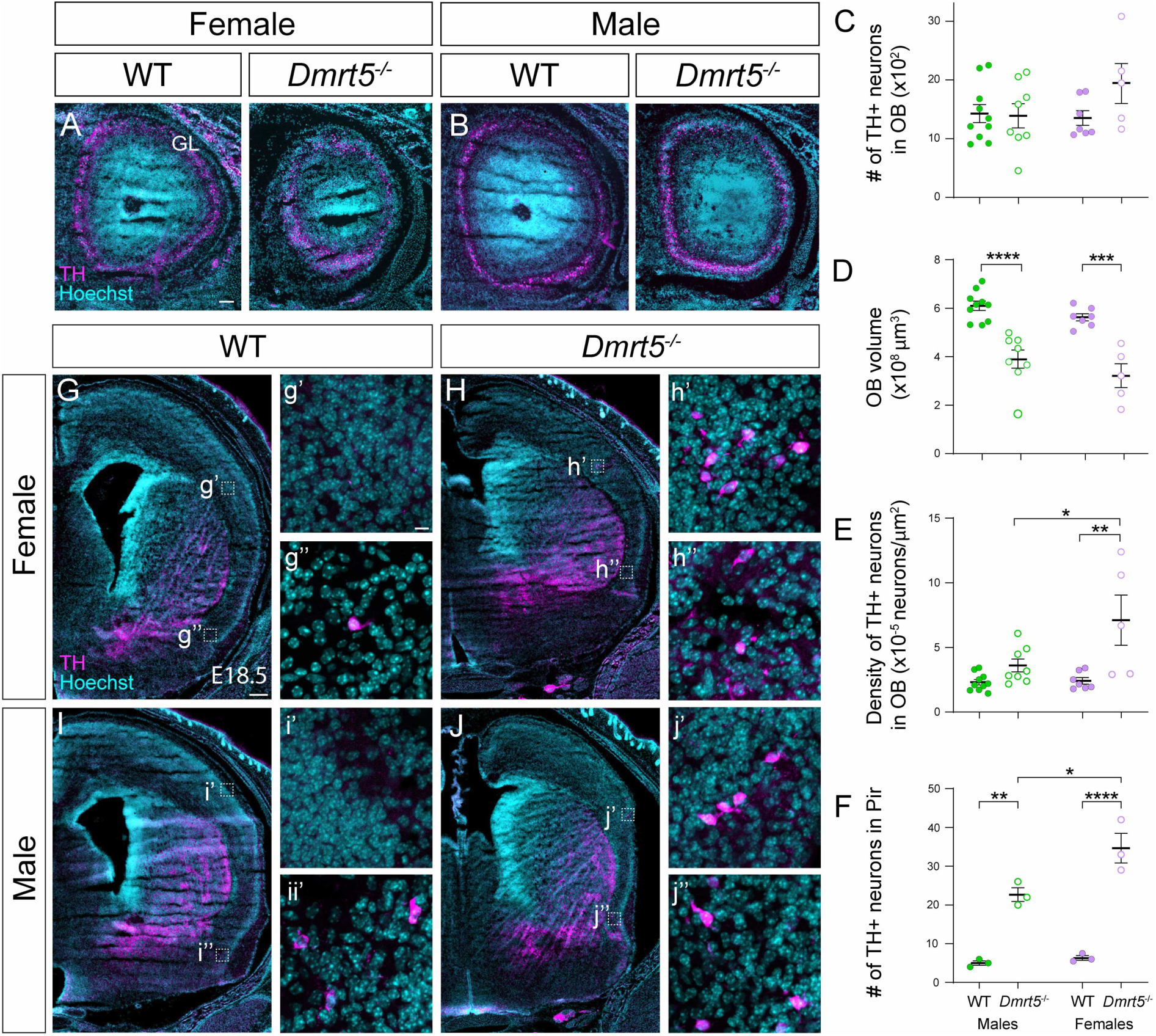
The number of TH neurons increases in *Dmrt5^-/-^*female embryos. **A, B)** TH antibody staining in OB slices of WTs and *Dmrt5^-/-^*in both sexes at E18.5. The GL layer is thicker in the mutants. Scale bar: 100 µm. **C-E)** Quantification of TH-OB neurons. The graphs represent quantifications expressed as mean±SEM from whole OBs in 3D-claired brain reconstructions**. C)** Total number of TH+ neurons, **D)** OB volume, **E)** Density of TH neurons (number of TH/Volume). **G-J)** TH IF in E18.5 brain slices in the Pir and neocortex of WTs and *Dmrt5^-/-^* in both sexes. Ectopic TH neurons are detected in *Dmrt5^-/-^* mutant neocortices in both sexes (**g’-j’**). Scale bars: 200 µm (G) and 10 µm (g’). **F)** There is a higher increase in TH-Pir neurons in *Dmrt5^-/-^* in female embryos (**g’’-j’’**). The graph represents mean±SEM from whole 3D-claired brain reconstructions. Each dot corresponds to a hemibrain. For each embryo, the two hemibrains were considered independently. Two-way ANOVA followed by Tukey multiple comparisons test: * p-value < 0.05, ** p-value < 0.01; *** p-value < 0.001; **** p-value < 0.0001.

Taking advantage of the TH staining in the clarified brains, we quantified the effect of *Dmrt5* in a group of TH+ neurons located within the Pir, at the boundary between Pir and the Tu (**Fig. 8G-8J**). These DA neurons, of unknown function, were PAX6+, which suggests their PSB origin (**Suppl. Fig. 5**). They have been previously reported to be derived from the PSB, where *Dmrt5* and *Dmrt3* are strongly expressed within the *Pax6* territory^45,76^. The number of Pir-TH in WT embryos is about the same in the two sexes. *Dmrt5*^-/-^ embryos presented a significant increase in Pir-TH neurons in both male and female embryos. Moreover, *Dmrt5*^-/-^ females presented significantly 1.55-fold more Pir-TH neurons than *Dmrt5*^-/-^ males (**Fig. 8F**). *Dmrt5* explained the majority of the variance in Pir-TH neuron numbers (82.78%, p < 0.0001 by two-way ANOVA), whereas sex (6.96%, p < 0.0148) and *Dmrt5*-sex interaction (4.45 %, p < 0.0384) contributed modestly, indicating that the interaction between *Dmrt5* and sex, although small, did exist.

In addition to the Pir-TH neurons, we found that in *Dmrt5^-/-^* embryos, TH/PAX6*+* cells appeared ectopically at the border of the neocortex and the Pir (**Fig. 8H**, **8J**). These neurons were never detected in WT brains and appeared in both sexes.

These results indicate an interaction between *Dmrt5* and sex in DA neurons from olfactory structures. These neurons originate from the PSB domain, where all DMA-*Dmrts* are expressed in early embryos (**Fig. 1C’-1E’**). Depletion of *Dmrt5* leads to the expansion of the PSB more dorsally by converting the identity of cortical progenitors to that of the PSB^28,45^. As a result, the persistent expression of PAX6 and an increase in OB-i were observed as they migrate to form the olfactory cortex. No effect was observed for OB-p neurons.

It would be interesting to check if there are any sexual differences in DMA-*Dmrt* expression levels in the PSB. The precise control of DMA-*Dmrts* dosage is crucial for cortical patterning, specifically in the PSB. Therefore, intrinsic differences in their expression between the sexes could have a significant effect.

## CONCLUDING REMARKS

The comprehensive analysis of *Dmrt* gene expression establishes the framework for understanding how this TF family contributes to mammalian brain development. We provide the first evidence of *Dmrt1* in the mouse brain and confirm robust expression of *Dmrt2* and *Dmrt6*, expanding their known roles beyond the gonadal and peripheral organs. The spatial and temporal distribution observed here suggests that *Dmrts* act in both unique domains to specify discrete neuronal populations and in overlapping regions where multiple family members may interact cooperatively.

Several regions emerged as particularly relevant. Multiple *Dmrts* co-localize in the PThE, a source of migrating neurons targeting brain regions such as the PO and hypothalamus, essential for innate behaviors^77^. In the MPall, most *Dmrts*, except *Dmrt2,* are co-expressed in progenitors that give rise to the hippocampus. Synergistic functions of *Dmrt3* and *Dmrt5* are well established here^45^, but the presence of *Dmrt1* and *Dmrt6* suggests additional roles yet to be defined. Interestingly, *Dmrt2*, undetected in hippocampal progenitors, appears in DG interneurons, involved in hippocampal plasticity^78^. The ventral midbrain also shows enriched *Dmrt* expression: *Dmrt5* in progenitors, *Dmrt4* in postmitotic DA neurons, and *Dmrt6* in distinct mantle regions. Such a spatial configuration raises the possibility of epistatic interactions regulating DA identity specification.

A key question in this study was whether these genes are sexually dimorphically expressed in the NS, given their evident dimorphism in gonads and invertebrate NS^3^. At the whole mRNA level, determined by ISH, we observed broadly similar expression between sexes, with only quantitative differences, such as higher *Dmrt2* expression levels in male CCx primordia^39^ or adult PO. Nonetheless, we did not delve into the intricate mechanisms of gene regulation, such as alternative splicing and post-translational regulation, which can also generate sex bias. The presence of splice variants predicted for *Dmrt2*, *Dmrt4,* and *Dmrt6*, and the precedent of sex-specific degradation through the DMA domain in *C. elegans*^79^, suggests additional layers of complexity. Functional analyses already suggest that *Dmrts* can buffer latent sex differences: knockdown of *Dmrt2* reveals male-biased effects on progenitor proliferation in the CCx^39^. Here we showed the absence of *Dmrt5* unmasks sex differences in Pir-TH neurons. Thus, *Dmrts* may serve as regulators of hidden sexual programs, only revealed under perturbation.

Our results also highlight roles that extend beyond embryogenesis. Most *Dmrts*, except *Dmrt1* and *Dmrt4*, persist into postnatal stages. The postnatal function of *Dmrts* has received limited attention, in part because most *Dmrt* null mutants do not survive past the perinatal period. Moreover, sex, as a biological variable, is frequently overlooked in neuroscience research^80^. Dmrt3 and Dmrt5 expression in SOX2+ progenitors in postnatal neurogenic niches strongly suggests they participate in the regulation of neural stem cell activity. Given known sex differences in hippocampal neurogenesis^81^, including estrogen-driven proliferation in females^82^ and testosterone-sensitive responses in males^83^, *Dmrts* could mediate intersections between hormonal signals and intrinsic cell-cycle regulation. The translational implications are notable. Human deletions at the *DMRT1–3* locus are associated with intellectual disability^84–86^, while *DMRTA2/DMRT5* mutations cause microcephaly in both humans and mice^48,87^, demonstrating evolutionary conservation. Similar mechanisms to those observed in the increase of TH neurons, where the interaction between *Dmrt5* deficiency and the sex of the organism modulates development, may operate in the hippocampus, hypothalamus, amygdala, and midbrain dopaminergic zones. This raises the possibility that dysregulated *DMRT* function contributes to sex differences in psychiatric symptomatology.

Looking ahead, several avenues stand out. First, advancing from transcriptional mapping to multi-omics integration—including proteomics and splicing analysis—will uncover hidden biases in *Dmrt* regulation. Second, conditional knockouts will be key for elucidating postnatal roles, given the perinatal lethality of null mutants. Finally, dissecting how *Dmrts* intersect with sex hormones and their receptors may unravel molecular links between developmental gene networks and sexual cues.

Despite limited overt transcriptional dimorphism, *Dmrts* exhibit complex spatial and temporal expression patterns, positioning them as critical regulators of brain development. Alternative splicing, protein regulation, and gene–gene interactions likely encode *latent* sex differences that shape neurogenesis, circuit formation, and disease susceptibility. By extending our understanding of DMRT biology into postnatal and human contexts, future research can uncover their contributions to cognitive functions and psychiatric disorders, with important implications for sex-specific vulnerabilities in brain disease.

**Suppl. Figure 1. Murine *Dmrt1* to *Dmrt6* loci.** Oligos used in *Dmrt2* RT-qPCR are indicated with red arrows. *Dmrt5^-/-^* mutation schematic has been adapted from Saulnier et al^44^.

**Suppl. Figure 2. *Dmrt6* expression in E18 and P7 brains.** *Dmrt6* ISH in male and female representative embryos is shown in the figure at E18.5 and P7. R, rostral; C, caudal.

**Suppl. Figure 3. *Dmrt2* expression in E18 brain.** *Dmrt2* ISH in male and female representative embryos is shown in the figure at E18.5. R, rostral; C, caudal.

**Suppl. Figure 4. *Dmrt5* expression in coronal and sagittal E18.5 brains.** *Dmrt5* ISH in male and female embryos. The first four rows depict a coronal view. R, rostral; C, caudal. The last four rows represent sagittal views of Dmrt5 ISH. L, lateral; M, medial.

**Suppl. Figure 5. PAX6 in TH neurons in *Dmrt5^-/-^* mutants.** PAX6 and TH double IF in WT and *Dmrt5^-/-^* embryos at E18.5, in females (**A**, **B**) and males (**C, D**). Ectopic TH neurons in the neocortex are PAX6+, white arrowhead (insets **a’’-d’’**).

## EXPERIMENTAL METHODS

### Animals and tissue preparation

Animals were housed and bred at the CBM animal facility and treated in accordance with European Communities Council Directive 86/609/EEC. All procedures were approved by the Bioethics Subcommittee of CSIC (Madrid, Spain) and the Comunidad de Madrid under the PROEX 286/19; RD 53/2013.

Animals used in this study include male and female C57BL/6JRccHsd (ENVIGO) and C57BL/6J (The Jackson Laboratory). Heterozygous mice B6J.Cg-*Sst^tm2.1(cre)Zjh^*/MwarJ (028864, The Jackson Laboratory) and B6.129P2-*Pval^btm1(cre)Arbr^*/J (017320, The Jackson Laboratory)^88^ were crossed with the reporter floxed mice *tdTomato* B6;129S6*-Gt(ROSA)26Sor^tm14(CAG-tdTomato)Hze^*/J (007914, The Jackson Laboratory).

For embryonic sex genotyping, tail tissue was collected and digested in 50 μl of MgCl_2_-free PCR reaction buffer (10013-4104, Biotools) supplemented with 400 μg/ml proteinase K (3115879001, Roche) at 55°C with shaking for 1-3 hours (h), followed by heat-inactivation of the proteinase K at 95°C for 5 min. The protocol of genomic DNA amplification was adapted from McFarlane *et al*^89^. The primers used were: *Sx*-Forward 5’-*GATGATTTGAGTGGAAATGTGAGGTA*-3’ and *Sx*-Reverse 5’-*CTTATGTTTATAGGCATGCACCATGTA*-3’.

The presence of a vaginal plug estimated embryonic day (E) 0.5 of pregnancy after overnight male mating. Embryos were collected from independent litters. For staining on slices, heads from embryos (E11.5, E12.5, E13.5, E14.5, E15.5, and E18.5) were fixed in 4% paraformaldehyde (PFA) (1.04005.1000, Merck) diluted in phosphate-buffered saline (PBS) for 2 h at room temperature (RT). Postnatal day (P) 4, P6 or P7 heads were fixed in 4% PFA overnight (o/n) at 4°C. Adult brains (P56) and adult noses (P71) were fixed in 4% PFA o/n at 4°C from animals previously perfused intracardially with sterile PBS followed by 4% PFA. After fixation, samples were washed several times in PBS. Adult noses were decalcified in 0.25 M EDTA for 3 days at 4°C and, after that, washed several times in PBS All samples were cryoprotected in 15%-30% sucrose (1.07651.1000, Merck) in PBS. Lastly, they were embedded in 7.5% gelatin (G2625, Sigma)/15% sucrose (1.07651.1000, Merck)/PBS and frozen at -80°C. Coronal and sagittal cryostat sections were produced at 16 μm thickness.

### *In situ* hybridization (ISH)

Tissue sections were hybridized with digoxigenin-labeled riboprobes as described in a previous work with minor modifications^90^. In the case of E12.5 embryos, the analysis was performed in coronal and sagittal planes on consecutive sections for at least three embryos of both sexes. In the case of postnatal tissue from P25 onward, sections were firstly treated with proteinase K (10 μg/ml; Merck, 3115879001) for 5 min, washed in PBS (pH 7.4), and post-fixed for 10 min in 4% PFA. For embryonic and neonatal tissues, the procedure began directly with post-fixation for 10 min in 4% PFA. After fixation, sections were washed in PBS (pH 7.4), acetylated, and washed in PBS with 0.1% Triton X-100. Then, sections were incubated for 1 h at RT with hybridization buffer [50% deionized formamide (Millipore, S4117), 1X salts (pH 7.5, 10 mM Tris, 200 mM NaCl, 5 mM NaH_2_PO_4_, 5 mM Na_2_HPO_4_, 5 mM EDTA), 1X Denhardt’s (Merck, D2532), 10% dextran sulphate (Merck, 4911), tRNA 1 mg/ml (Merck, R6625)] in a humified chamber with 5X SSC (pH 4.5, 750 mM sodium citrate, 75 mM NaCl) and 50% deionized formamide (Millipore, S4117). Then, sections were hybridized o/n at 72°C with riboprobes (1:1000 in hybridization buffer). After hybridization, sections were washed for 90 min in 0.2X SSC at 72°C, and blocked in 2% blocking solution (Roche, 11096176001) in 1X MABT [pH 7.5, 100 mM maleic acid, 150 mM NaCl, 0.1% Tween-20 (Merck, P1379)] for 1h at RT. After blocking, sections were incubated o/n at RT with anti-DIG antibody conjugated with the alkaline phosphatase (1:5000, Roche, 11093274910) in MABT. After extensive washing in TBST (100 mM Tris, 150 mM NaCl, 0.1% Tween-20) at pH 7.5 and then in TBST at pH 9.5, the alkaline phosphatase activity was detected at RT by using 100 μg/ml 4-nitro blue tetrazolium chloride (Roche, 11383213001) and 175 μg/ml 5-bromo-4-chloro-3-indolyl phosphate *p*-toluidine salt (Roche, 11383221001) in NTMT (pH 9.5, 100 mM Tris, 100 mM NaCl, 50 mM MgCl_2_, 0.1% Tween-20). Sections were mounted in Mowiol (Merck, 81381).

Riboprobes were generated in this study *de novo* for *Dmrt6*, *Dmrt7, and GlyT2* genes (**Suppl Fig. 1**; **Table 3**). The oligos used to isolate the cDNA sequences were the following: for *Dmrt6* (5’-*ATGCTTCGCGCCCCCAAGTG*-3’ and 5’-*TGCTCCTGGGGAGAGTCCTG*-3’); for *Dmrt7* (5’-*TGCTCTCCACCACTGTTCTG*-3’ and 5’-*GCTACAAGAATAGCCACCCTCAG*-3’); for *Esr2 (5’-GCCGACTTCGCAAGTGTTAC*-3’ and 5’-*GCTTCGAGGGTACAAGTCCTCA*-3’), and for *GlyT2* (5’-*ACAGCGCACAGGCCAATCC*-3’ and 5’-*GTTCGGATAGGCAGTCATGCAG*-3’). The rest of the riboprobes have been published before and kindly donated by Prof. Eric Bellefroid for *Dmrt1*, *Dmrt3*, *and Dmrt5*; Dr. Nuria Flames for *Vglut2*^91^ and *gad65,* Dr. Takahiko Sato for *Dmrt2,* and Dr. Dajiro Konno for *Dmrt4* (**Suppl Fig. 1**; **Table 3**).

**Supplementary Table 1.** Riboprobes used in *in situ* hybridization.

| Gene | Probe sequence information | Reference |
| --- | --- | --- |
| *Dmrt1* | 1299 bp (covers cDNA) | Gift from Prof. Eric Bellefroid |
| *Dmrt2* | 1686 bp (covers cDNA) | Ref 44 |
| *Dmrt3* | 1432 bp (covers cDNA) | Ref 53 |
| *Dmrt4* | 1470 bp (covers cDNA, except ATG) | Ref 26 |
| *Dmrt5* | 891 bp (covers 3rd exón + 3'UTR) | Ref 53 |
| *Dmrt6* | 1070 bp (covers cDNA) | This study |
| *Dmrt7* | 1265 bp (within exon 1-3'UTR) | This study |
| *Esr2* | 903 bp (within exon 3- exon 8) | This study |

### Immunofluorescent staining (IF)

IF was performed following standard protocols. Brain sections were washed with PBS with 0.1% Triton X-100 (T9284, Sigma-Aldrich) (PBST) and blocked for 1h in blocking solution [PBST and 1% of bovine serum albumin (BSA) (A2153-100G, Sigma)]. Primary antibodies were diluted in a 1/10 blocking solution and incubated o/n at 4°C; the corresponding secondary antibodies were incubated at 1:1000 for 2h at RT. Hoechst 33342 (1/1000) (H1399, Invitrogen™) was used for cell nuclei staining. For double ISH or FISH and IF staining, ISH was performed before IF, as previously described^90^.

**Supplementary Table 2.** Primary antibodies.

| Antibody (epitope) | Host specie | Reference | Dilution |
| --- | --- | --- | --- |
| βIII Tubulin (Tuj1) | Mouse | Promega (G7121) | 1:500 |
| BRN3A (POU4F1) | Goat | Santa Cruz Biotechnology (sc-31984) | 1:2000 |
| DMRT3 | Guinea Pig | Gift from Irene Miguel-Aliaga | 1:400 |
| DMRT5 | Rabbit | Ref 59 | 1:2000 |
| LHX3 | Rabbit | Gift from Tom Jessell lab (Columbia University) | 1:200 |
| OMP | Goat | FUJIFILM Wako (019-22291) | 1:1000 |
| Reelin | Mouse | Millipore (MAB5364) | 1:300 |
| SOX2 | Goat | R&D (AF2018) | 1:50-250 |
| TH | Rabbit | PelFreez (P4010) | 1:400 |
|  | Mouse | Merck (T1299) | 1:600 |

### Imaging

Chromogenic ISH images were acquired on a Leica DM5000B vertical microscope equipped with a Leica DFC500 color camera. IF images were acquired on the previous Olympus SpinSR10 confocal, a Leica DM5000B vertical microscope fitted with a Leica DFC350 FX monochrome camera, or a Zeiss LSM900 confocal coupled to a vertical Axio Imager 2 microscope.

### Immunolabeling-enabled three-dimensional imaging of solvent-cleared organ plus (iDISCO+)

Tissue preparation and whole-mount immunolabeling were performed using the iDISCO+ protocol, optimized for embryonic tissues^92^. E18.5 embryos were collected in ice-cold Leibovitz L-15 medium or PBS, and allowed to drain blood for 5 minutes on ice before fixation in 4% PFA at 4°C o/n with agitation.

Fixed samples were pretreated with methanol for tissue dehydration and bleaching. This involved a graded methanol series (20–100%, 1hr each at RT with shaking) followed by o/n incubation in a 66% dichloromethane (DCM) (5.89581, Sigma-Aldrich)/33% methanol (2:1 v/v; 1.06009, Supelco) solution with shaking. After additional washes in 100% methanol, samples were bleached in freshly prepared 5% hydrogen peroxide (H₂O₂)/methanol o/n at 4°C. Following rehydration through a reverse methanol gradient to PBS, samples were washed twice in PTx.2 buffer (2% Triton X-100/PBS) for 1h each at RT with shaking.

Pre-treated samples are permeabilized using a solution containing 0.3 g/mL Glycine and 20% DMSO (472301, Sigma-Aldrich) in PTx.2 at 37°C with shaking for up to two days. Samples were then blocked using a 6% Donkey Serum (017-000-121, Jackson Immunoresearch) and 10% DMSO/PTx.2 blocking buffer at 37°C with shaking for up to two days. Samples were incubated with the primary antibody in PTwH (10-50 µg/ml Heparin sodium salt in 0.2% Tween20/PBS) supplemented with 5% DMSO and 3% donkey serum for three (E18.5 brains out of skull) or four (E18.5 embryos head) days at 37°C with shaking. Rabbit anti-TH at 1:1000 (Merck; AB152) was used; For DMRT3, a guinea pig anti-DMRT3 at 1:400 (a gift from Irene Miguel-Aliaga) was applied. Subsequently, samples were extensively washed in PTwH and left in PTwH o/n at 4°C before proceeding with secondary antibody incubation. The secondary antibodies used were Donkey anti-Rabbit IgG (H+L) Highly Cross-Adsorbed Secondary Antibody, Alexa Fluor 647 (at 1:500; Jackson ImmunoResearch; 711-605-152. Secondary antibodies were applied under similar conditions to primary antibodies for three or four days. Antibody solutions were filtered through a 0.2 µm membrane or centrifuged at 20,000 × g for 10 min before use to minimize the formation of precipitate. After several washes in PTwH, clearing was achieved by dehydrating samples through a methanol gradient followed by incubation in DCM/Methanol (2:1 v/v) for 3h, and two additional 15-minute incubations in 100% DCM. Final optical clearing was performed in dibenzyl ether (DBE) (108014, Sigma-Aldrich), with samples fully immersed in DBE-filled tubes to minimize oxidative damage.

Cleared samples were placed in an imaging reservoir made of 100% quartz filled with ethyl cinnamate (ECi) (W243000, Sigma-Aldrich) and illuminated from the side by a laser light using a LaVision Biotec Ultramicroscope II (Miltenyi Biotec, Germany)/ImspectorPro (LaVision BioTec) software. For DMRT3, ECi was used to fill the sample chamber, and cleared samples were fixed to the sample holder using UV adhesive (Bondic, BC5000) for 3D imaging using a ZEISS Lightsheet 7 (Carl Zeiss, Germany)/Zeiss Zen Black (Carl Zeiss, Germany) software. 3D images and their subsequent processing were carried out using Imaris x64 software (version 9.3.1, Bitplane). Zeiss Arivis Pro (version 4.3.0, Carl Zeiss) for DMRT3 DG stainings. The different volume reconstructions (OB and Pir) were generated manually by creating a mask using the “surface tool” and they were pseudo-colored. The TH+ cell number was obtained using the “spots tool”, which was applied to each of the regions under study, and was conducted on both brain hemispheres to enable the detection of potential interhemispheric differences in marker expression, structural organization, or cellular distribution. This approach enhances the sensitivity of the analysis to subtle anatomical or molecular differences that may be obscured in unilateral assessments. It ensures that any hemispheric bias in staining, labeling, or tissue handling does not compromise the accuracy of the results.

### RNA isolation and qRT-PCR

Total RNA was isolated using NZY Total RNA Isolation kit (MB13402, Nzytech) following the manufacturer’s instructions. cDNA was obtained from 1 μg of total RNA with First-Strand cDNA Synthesis kit (27-9261-01, GE) in a 15 μl reaction volume. qPCR was performed using GoTaq® qPCR Master Mix (A6002, Promega) according to the manufacturer’s protocol in a CFX-384 Touch Real-Time PCR Detection System (Bio-Rad). The following gene-specific primer pairs were used: *Dmrt2-*Forward 5’-*CCTGAGCCTGGTTCTTG*-3’, *Dmrt2*-Reverse 5’-*CAGTGGAGTCCCACAGCTATC*-3’, *Gus*-Forward 5’-*AGCCGCTACGGGAGTCG*-3’, and *Gus*-Reverse 5’-GCTGCTTCTTGGGTGATGTCA-3’. *Dmrt2* and *GFP* expression levels were quantified in triplicate and normalized to those of Gus. The data were analyzed using the comparative Ct method.

### Analysis of the MOE in *Dmrt5^-/-^* null mutants

16-micron coronal cryostat sections of the MOE were prepared from *Dmrt5^-/-^* null mutant and WT E14.5 embryos of both sexes. Sections were immunostained with primary anti-OMP antibody, followed by incubation with Alexa Fluor 555-conjugated secondary antibody (1:1000). Nuclei were counterstained with Hoechst 3334 (1:1000). For each sample, a single section representing the central region along the anteroposterior axis of the unilateral MOE was selected for analysis. Quantification of OMP+ cells and measurement of MOE area were performed on these selected sections.

To determine the number of OMP+ cells, immunostained sections were imaged using an Olympus SpinSR10 spinning disk confocal system mounted on an IX83 inverted microscope. For complete coverage of the unilateral MOE, 1–2 images were acquired per sample using a 20x objective. Each image consisted of a Z-stack comprising 51 optical sections, which were subsequently Z-projected to generate a single image. OMP+ cells were manually counted on the Z-projected images using the Cell Counter plugin in Fiji (ImageJ). OMP+ cells located outside the thickness of the epithelium were not counted.

To measure MOE area, each sample was imaged with Hoechst nuclear staining. Images were acquired using a Leica DM5000B upright microscope equipped with a Leica DFC350 FX monochrome camera and a 10x objective. The unilateral MOE area was delineated and measured using the freehand selection tool in Fiji.

OMP+ cell density for each unilateral MOE was calculated by dividing the number of OMP+ cells by the corresponding MOE area.

### Statistical analyses

The researchers were blinded to the sex and genotype at the time of analysis. The number of quantified animals is indicated in each plot; a minimum of three mice were included. The normality of the distribution was tested using the Shapiro-Wilk test in all cases. Each individual test is indicated in the figure legend of the corresponding experiments. Statistically significant differences were considered and represented by p-value: *p < 0.05, **p < 0.01, and *** p < 0.001. Statistical analyses were conducted using GraphPad Prism 9 and 8.0.2.

## ACKNOWLEDGMENTS

We thank Prof. Eric Bellefroid for *Dmrt* probes and *Dmrt5^-/-^* null mutant mice. Prof. Irene Miguel-Aliaga for DMRT3 antibody. Dr. Nuria Flames and Dr. Takahiko Sato for *in situ* probes. The Advanced Light Microscopy and Animal facilities at the CBM (CSIC-UAM). We also thank Prof. Paola Bovolenta, Dr. Leonardo Beccari, and lab members for discussion and advice. Dr. Chen Wang and Dr. Eduardo Leyva for critical reading.

## STATEMENTS & DECLARATIONS

Ethics approval and consent to participate are not applicable.

Written informed consent for publication was obtained from all participants included in the study.

This published article and its supplementary information files include all data generated or analyzed during this study. In situ raw data are available at https://www.informatics.jax.org/reference/J:364871. MGI Direct Data Submission to Mouse Genome Database (MGD). The Jackson Laboratory, Bar Harbor, Maine. World Wide Web (URL: http://www.informatics.jax.org).

Grant PGC2018-101751-A-100 and PID2021-127235NB-I00 funded by MCIN/AEI/10.13039/501100011033, and ERDF, “A way of making Europe”.

RC-N holds an FPU from the Spanish MCIN, FPU19/02352; AB-S holds an FPI fellowship from the Spanish MCIN, PRE19-089366; RTC holds an FPI from the CAM, PIPF-2022/SAL-GL-24909, cofounded by ESF. LC-C holds a JAE from CSIC, JAEINT24_EX_0840 SJ was supported by Grant PID2020-113878RB-I00/AEI/10.

ES-S. was supported by Grant RYC-2016-20537 funded by MCIN/AEI/10.13039/501100011033 and ESF, “Investing in your future”.

All authors declare they have no financial interests.

## AUTHORS’ CONTRIBUTIONS AND INFORMATION

ES-S conceived and designed the study and supervised the project. ES-S, RC-N, AB-S, and RTC developed the theoretical framework. All authors discussed the results, ES-S, RC-N contributed to the writing of the manuscript.

RC-N performed the ISH for *Dmrt3*-*Dmrt7*, as well as their anatomical identification. Additionally, RC-N conducted functional experiments to investigate the role of *Dmrt5* in the olfactory system. AB-S performed all ISH for *Dmrt2*. RTC performed iDISCO stainings for DMRT3 in the DG. V-O did the DMRT5 double stainings in the DG of postnatal animals. LC-C did *Dmrt1* and *Dmrt7* ISH. SA-B, MPM-V y SJ contributed with TH iDISCO staining in wildtype and *Dmrt5^-/-^* brains, including quantifications between sexes. All authors approved the final version.

## Notes

https://www.informatics.jax.org/reference/J:364871

### Competing Interest Statement

The authors have declared no competing interest.

