## Supplementary figures and images for "Expression Atlas Of *Dmrt* Genes Across Sex And Development – Functional Insights From The Mouse Olfactory System"

### Supplemental Figures

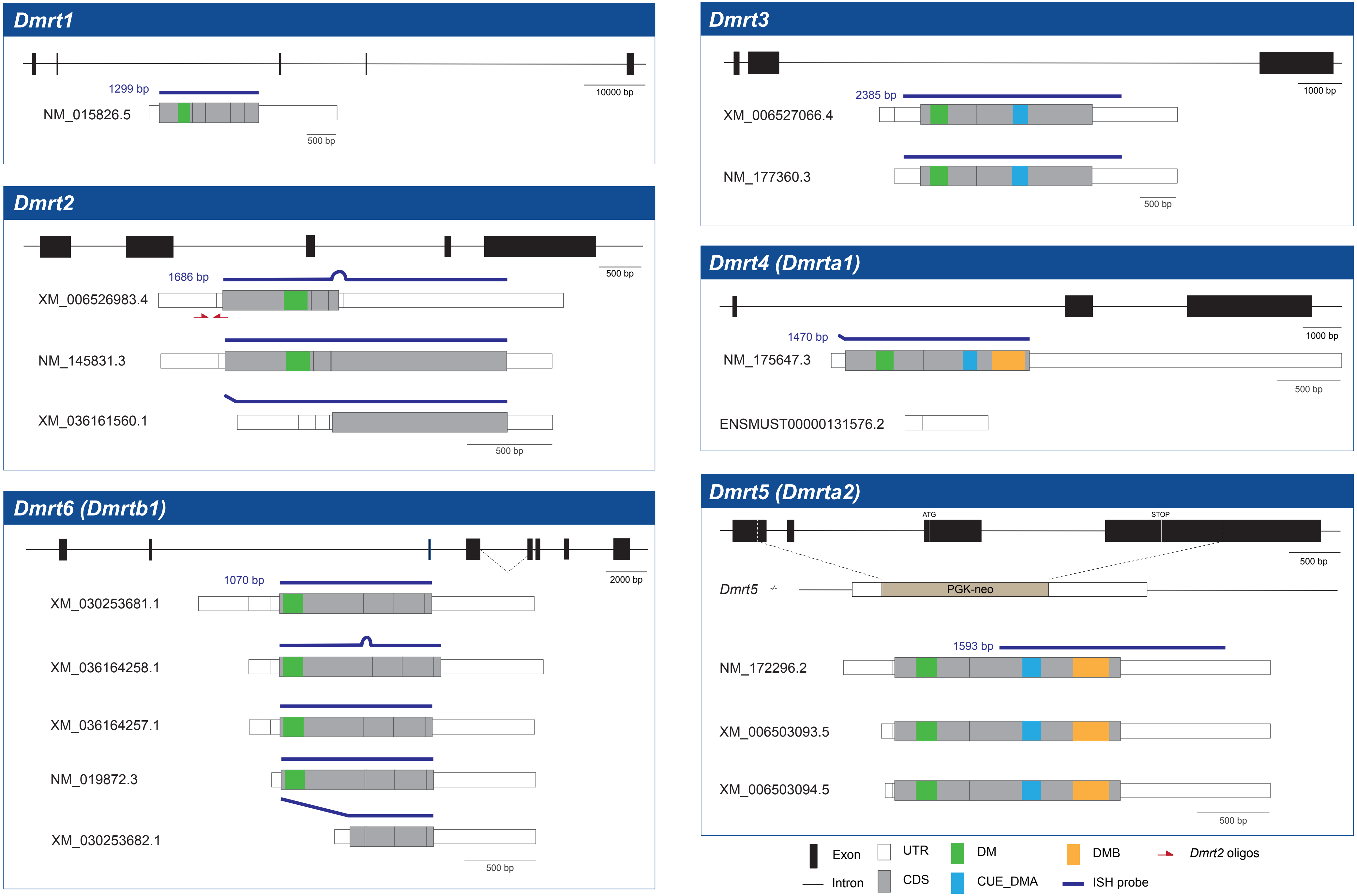

Suppl. Fig. 2

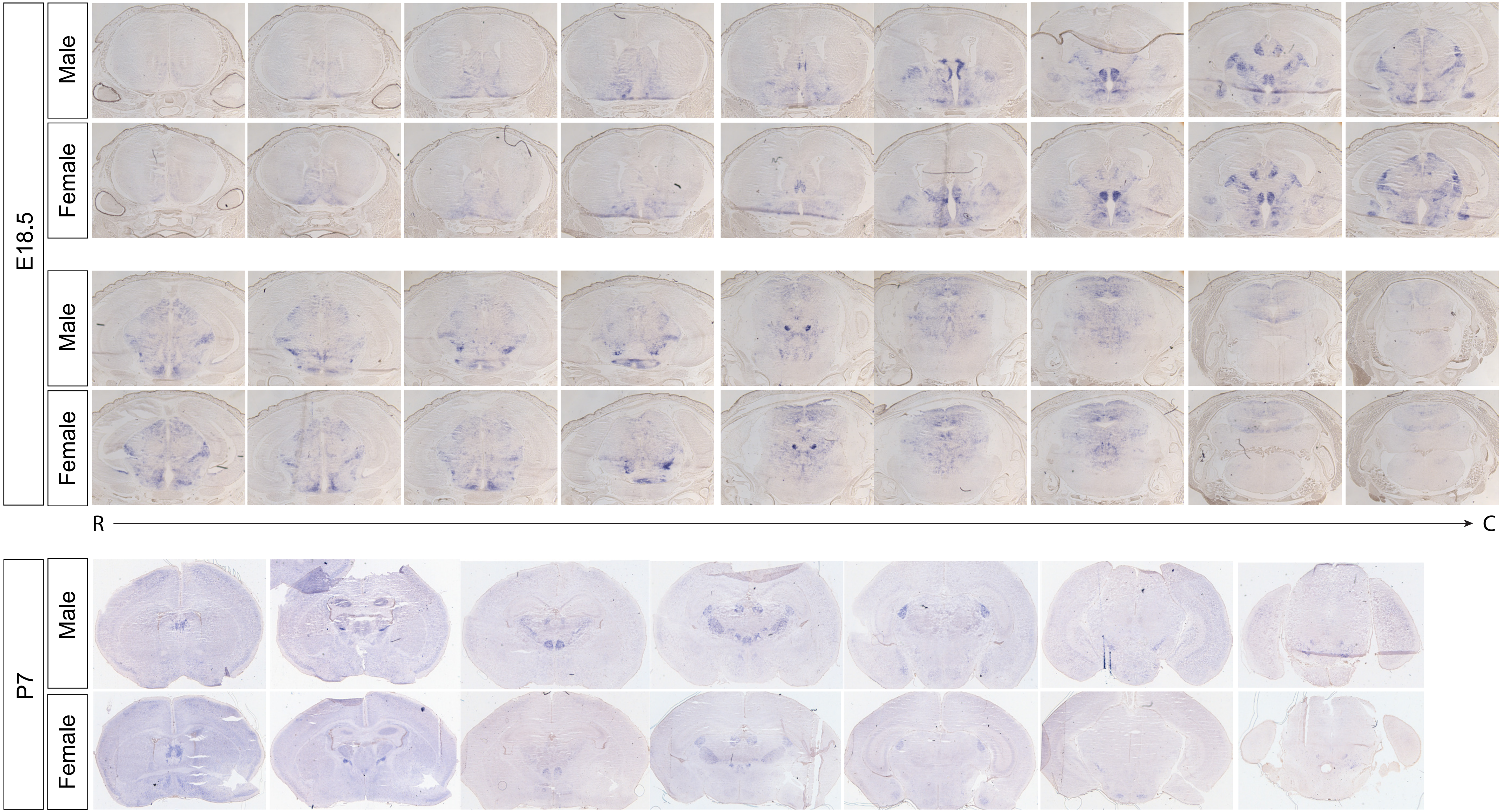

Suppl. Fig. 3

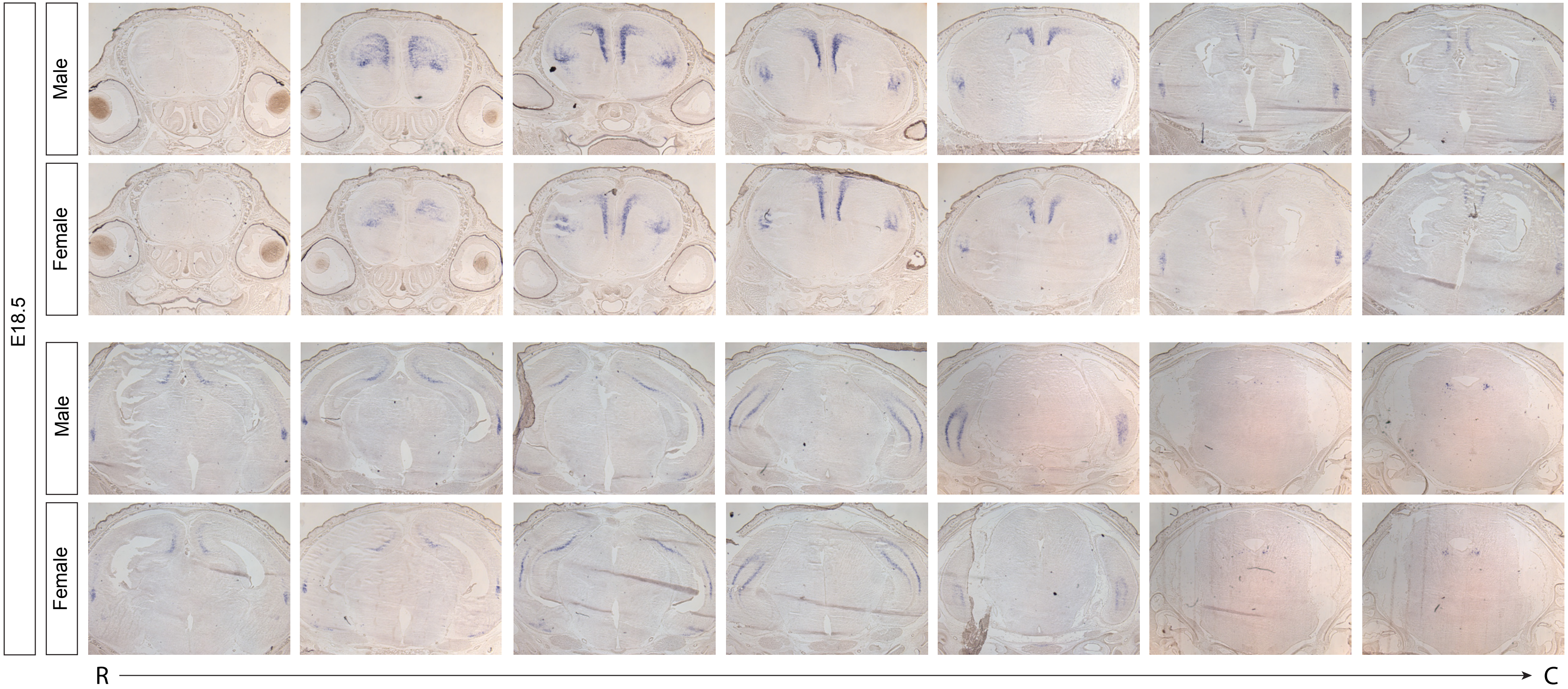

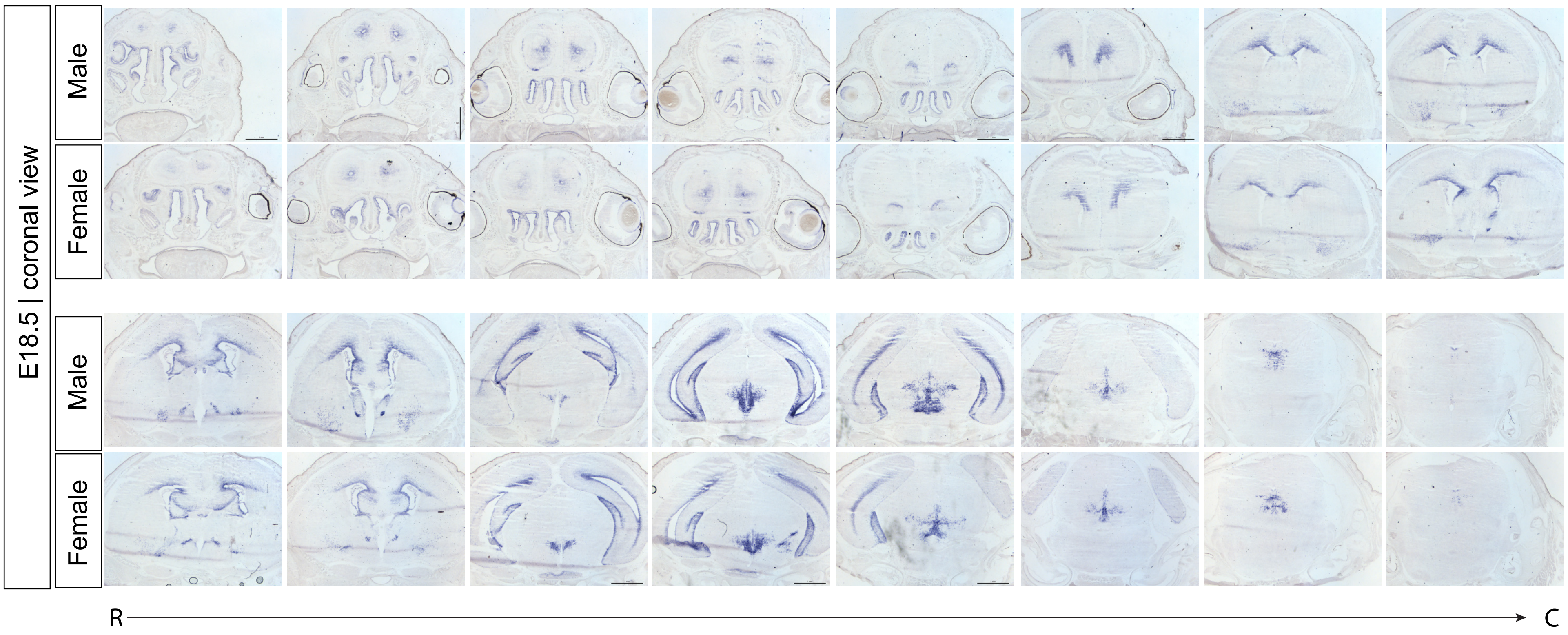

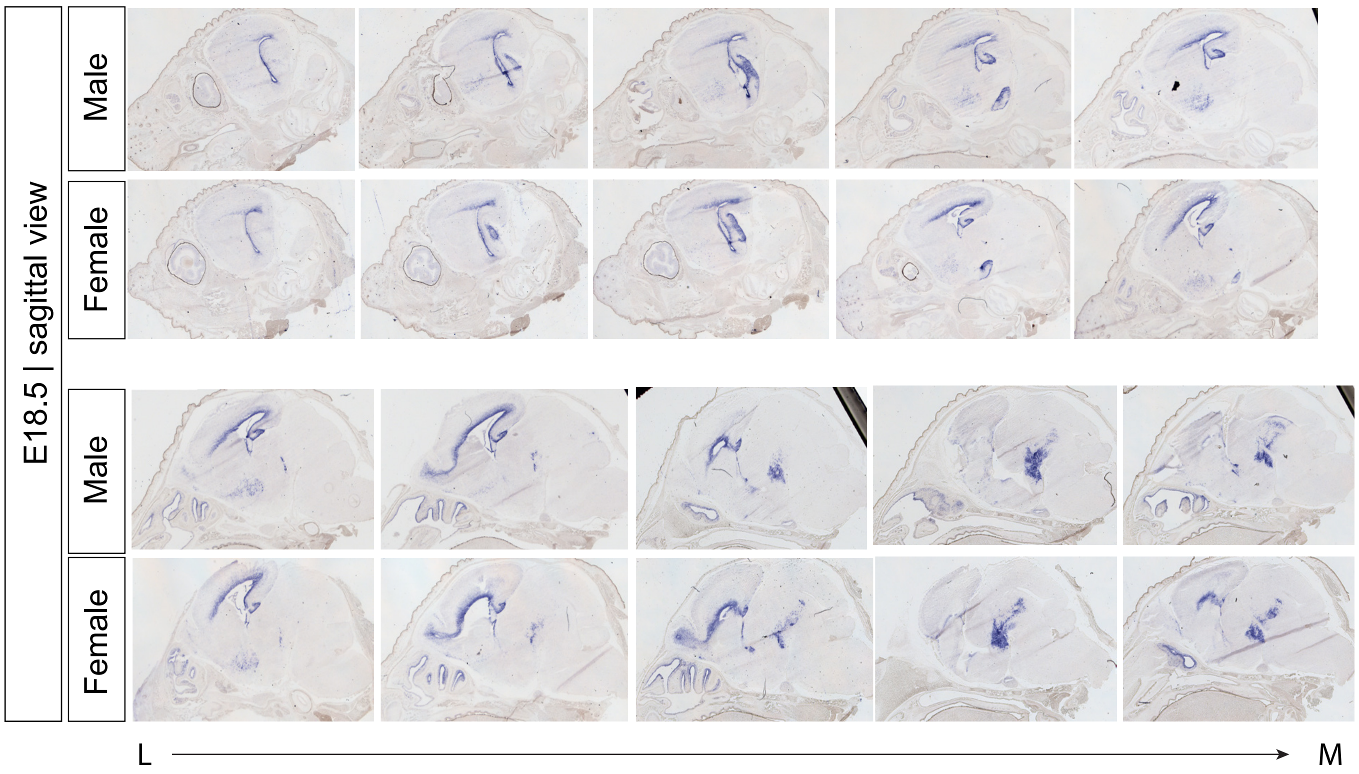

Suppl. Fig. 5

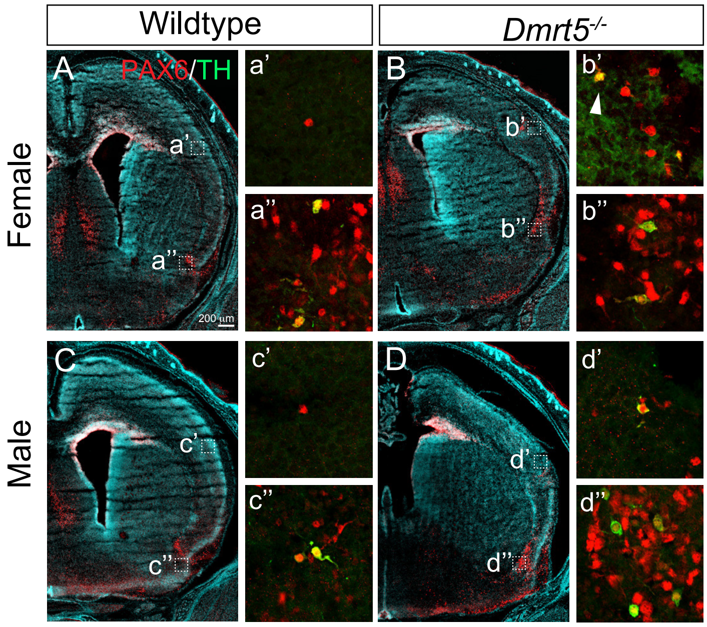
